# Cheminformatics identification of phenolics as modulators of penicillin binding protein (PBP) 2x of *Streptococcus pneumoniae* towards interventive antibacterial therapy

**DOI:** 10.1101/2023.10.02.560627

**Authors:** Jamiu Olaseni Aribisala, Nosipho Wendy S’thebe, Saheed Sabiu

**Author notes:** Corresponding author: Saheed Sabiu.

## Abstract

Infections caused by multidrug-resistant *Streptococcus pneumoniae* remain the leading cause of pneumonia-related deaths in children < 5 years globally, and mutations in penicillin-binding protein (PBP) 2x have been identified as the major cause of resistance in the organism to beta-lactams. Thus, the development of new modulators with enhanced binding of PBP2x is highly encouraged. In this study, phenolics, due to their reported antibacterial activities, were screened against the active site of PBP2x using structure-based pharmacophore and molecular docking techniques, and the ability of the top-hit phenolics to inhibit the active and allosteric sites of PBP2x was refined through 120 ns molecular dynamic simulation. Except for gallocatechin gallate and lysidicichin, respectively, at the active and allosteric sites of PBP2x, the top-hit phenolics had higher negative binding free energy (ΔG_bind_) than amoxicillin [active site (−19.23 kcal/mol), allosteric site (−33.75 Kcal/mol)]. Although silicristin had the best broad-spectrum effects at the active (−38.41 kcal/mol) and allosteric (−50.54 kcal/mol) sites of PBP2x, the high thermodynamic entropy (4.90 Å) of the resulting complex might suggest the need for its possible structural refinement for enhanced potency. Interestingly, silicristin had a predicted synthetic feasibility score of < 5 and quantum calculations using the DFT B3LYP/6-31G+ (dp) revealed that silicristin is less stable and more reactive than amoxicillin. These findings point to the possible benefits of the top-hit phenolics, and most especially silicristin, in the direct and synergistic treatment of infections caused by *S. pneumoniae*. Accordingly, silicristin is currently the subject of further confirmatory *in vitro* research.

## 1.0 Introduction

The incidence of antibiotic-resistant *Streptococcus pneumoniae* has been on the increase since its first report in the 1960s [1]. Globally, *S.* pneumoniae is the leading cause of community-acquired pneumonia, bacteremia, meningitis, and otitis media [2] and in children < 5 years of age, the pathogen remains the leading cause of pneumonia-related deaths [3–5]. With more than half of *S. pneumoniae* isolates frequently showing reduced susceptibility to beta-lactam antibiotics [6,7], the most frequently administered antibiotics for pneumococcal infections, treatment of the diseases has become problematic, necessitating the need for the discovery and development of novel drug candidates with an alternative mode of binding of key therapeutic target of the organism [6]. Penicillin-binding proteins (PBPs) are the typical target enzymes for beta-lactams and are engaged in the late phases of peptidoglycan biosynthesis, where their essential function involve transpeptidation of two muropeptides [8]. Mutation of important PBPs is a resistance mechanism which has been described in various bacteria species [9, 10]. The mutation entails modifying key PBPs such that interaction with beta-lactams occurs at considerably higher antibiotic doses than with PBPs from susceptible strains, and therefore the biological activity of the antibiotics is significantly diminished [8]. Thus, identifying improved modulators against critical PBPs of *S. pneumoniae* may aid in the battle against infections caused by multidrug resistant *S. pneumoniae*.

*S. pneumoniae* possesses six PBPs, including PBP1a, PBP1b, and PB2a, which are Class A high-molecular-mass PBPs (hmm-PBPs), PBP2x and PBP2b (Class B hmm-PBPs), and PBP3, a D, D-carboxypeptidase low-molecular-mass PBP (lmm-PBP) [11]. Antibiotic resistance in *S. pneumoniae* has been linked to all six PBPs [9]. However, only the class B hmm-PBPs (PBP2x and PBP2b) are key beta-lactam targets, and thus far, only PBP2x has been characterized as a major PBP target in clinical isolates since PBP2x genes that express low-affinity variants of *S. pneumoniae* can transfer beta-lactam resistance to susceptible strains [12]. Also, resistance due to mutations in PBP 2b and 1a of *S. pneumoniae* has been shown to occur in strains that already have a low affinity PBP2x, suggesting PBP2x as the most essential PBP in *S. pneumoniae* that is responsible for resistance [12]. Beta-lactams interact with PBP2x by acylating the active (actv) site serine residue located in the first of three conserved motifs (S*XXK, SXN, and KS/TG) of PBP2x, and mutations in this region have been attributed to resistance, which is the primary resistance mechanism for beta-lactam antibiotics in *S. pneumoniae* (Tsui et al., 2014). Interestingly, a PASTA domain located adjacent to the actv site (27 Å), whose function has been well-deciphered over the years, has in recent times been demonstrated to actv as an allosteric (allo) site in the generation of promising peptidoglycan and thus, when appropriately bound, could aid beta-lactam entry into the actv site [12, 13]. Bernardo-Garca *et al*. [12] showed that the complexation of a pentapeptide strain at the PASTA domains prompts another pentapeptide strand in the same nascent peptidoglycan strand to enter the actv site for the turnover events. Thus, the PASTA domain of PBP2x could serve as a good therapeutic target for identifying drug candidates that could actv in synergy with beta-lactams [14], and in this study, both the actv site and the allo site of PBP2x were employed in the identification of drug candidates against infections caused by *S. pneumoniae*.

Plant and plant-derived metabolites have long been used in the treatment of human diseases. Interestinling the increased interest in plant-based antimicrobials in recent years, particularly phenolic compounds, has resulted in the identification of numerous phenolic candidates demonstrating promising pharmacological properties [15, 16]. Phenolics ability to cause membrane damage, bind to virulence factors such as PBPs, and cause the suppression of bacteria biofilm formation, among other properties, has been implicated in its ability to inhibit both Gram-positive and Gram-negative microorganisms [17, 18]. The characteristic hydroxyl groups attached to the benzene rings of phenolics have been shown to enhance strong protein binding while increasing affinity for cytoplasmic membranes [16]. Alongside their direct antibacterial activity against microorganisms, several phenolics have been shown to work in synergy with conventional antibiotics, an approach that is encouraged in the treatment of infections caused by multi-drug-resistant bacteria [19]. Several phenolics have been identified from various plant parts, with repositories in several databases [20]. Repurposing phenolics could have the potential to reduce the time and costs to identify potent antimicrobials against PBP2x of *S. pneumoniae* [20]. Thus, in this study, all the currently known phenolics were repurposed against PBP2x by compiling phenolics that could interact with the protein from the ZINCPharmer database using structural-based pharmacophore and molecular docking techniques. The limitations of virtual screening techniques relating to lack of energy refinement and assessment of stability were addressed using molecular dynamic simulation and mechanics/GB surface area (MMGBSA) approaches. This was carried out to identify potentially novel and actv antibacterial candidates that can treat or act in synergy with conventional antibiotics in the treatment of infections caused by *S. pneumoniae*.

## 2.0 Results and Discussion

### 2.1 Screening of phenolics at the actv site of PBP2x using structure-based pharmacophore and molecular docking

Following the evaluation of the binding positioning and interaction of the 1555 phenolics obtained via structure-based pharmacophore at the actv site of PBP2x using the molecular docking approach, the top 20 phenolics had higher docking scores than all the beta-lactam antibiotics used as standards. The top 20 phenolics had docking scores ranging between −8.4 and −9.9 kcal/mol, with ZINC03793048 having the highest docking score, while the beta-lactams had docking scores between −7.1 and −8.0 kcal/mol, with amoxicillin having the highest docking score (Figure 1), and hence it was selected as a standard for further analysis. This observation suggests the probable better interactions and orientation of the top-hit compounds at the actv site of PBP2x relative to the standards. However, as virtual screening often produces inactive molecules in the preclinical and clinical stages of drug development [21], the identified top 20 compounds were further screened cheminformatically to identify the top hit phenolics that will be pharmacokinetically-friendly, have drug-like properties, and have a synthetic score that can enable structural alterations for enhanced druggability (Figure 1). Using these approaches, the top hit phenolics obtained and the standards passed the Lipinski rule of 5 (LRo5), which serves as a pioneering physicochemical filter, relating the physicochemical parameters of compounds to their pharmacokinetic properties and oral bioavailability [22]. The LRo5 limits the molecular weight, hydrogen donor, hydrogen acceptor, and octanol coefficient of a bioactive drug to 500 g/mol, 5, 10, and 5, respectively (Figure 1). Also, the top-hit phenolics and the standards had a synthetic score of < 5 (Figure 2) that would encourage structural alteration to improve potency [23]. GCG (ZINC03870413) had the lowest synthetic score (4.20) among the top-hit phenolics, with a value that compares well with amoxicillin (4.17), suggesting that the compound will be the most susceptible to structural modification for enhanced druggability. Other predicted pharmacokinetic properties (Table 1), such as the bioavailability scores, solubility, and gastrointestinal tract absorption of the top-hit phenolics, all point to the drug-like properties of the compounds [24], as they all have characteristics that compare well with amoxicillin. The top-hit phenolics toxicity profiles suggest that they are non-inhibitors of most of the cytochrome P450 (CYP), an important isoenzyme in drug metabolism [25]. GCG and amoxicillin are non-inhibitors of all the isoenzymes, suggesting that they would not result in drug-drug interactions (DDI) when co-administered with phase I- and phase II-dependent drugs. However, other top hit phenolics inhibit CYP3A4, suggesting their potential ability to cause DDI with drugs that are metabolized by the isoenzyme. This observation pinpoints to the need for further structural alteration to improve the toxicity profile of the compounds, more so as some of the compounds [silicristin, lysidicichin, and ECMG] were also predicted to be active for immunotoxicity (Table 1). However, ECBE, GCG, and amoxicillin are predicted to be non-active for all the toxicity endpoints. Remarkably, all the top-hit phenolics belong to either toxicity class 4 or 5 and can be deployed as medications [22, 24]. Generally, the top-hit phenolics in this study have pharmacokinetic, synthetic, and drug-like properties that suggest them as potential drug candidates that can be structurally modified for enhanced potency.

**Figure 1:**
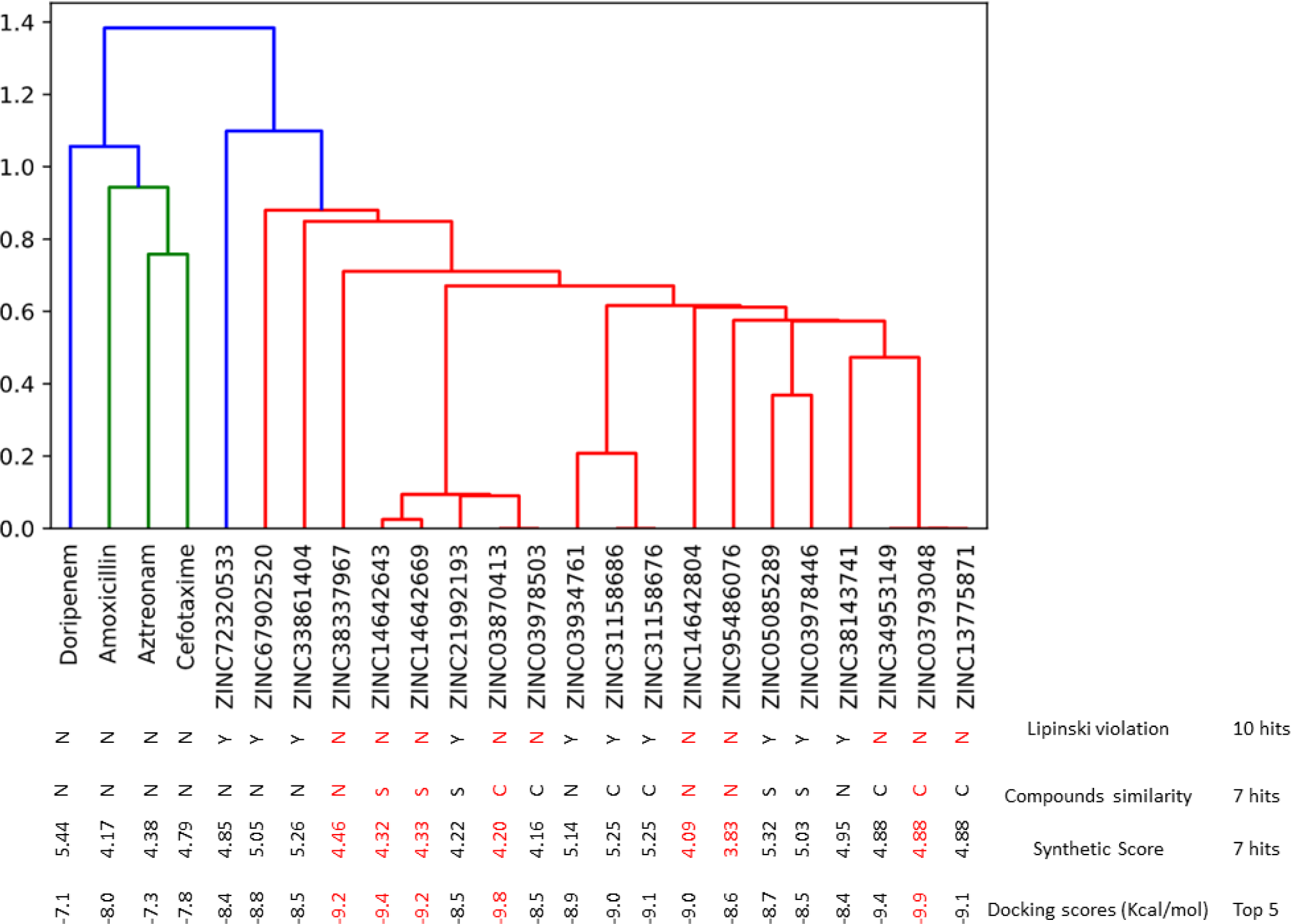
Cheminformatics and docking scores ranking of the top-hit phenolics. The same colour and cluster signifies more similarity. Except for ZINC72320533, all top twenty phenolics had the same colour meaning they are more similar than the beta-lactams with different colour. Compounds of the same clusters with zero similarity scores are conformers and in ranking the top-hit, only one (compounds that fulfilled the Lo5 and has the highest docking score) from a set of conformers are chosen. The top-hit phenolics all had a synthetic sore of < 5 and possess the resorcinol structure as a common moiety.

**Figure 2:**
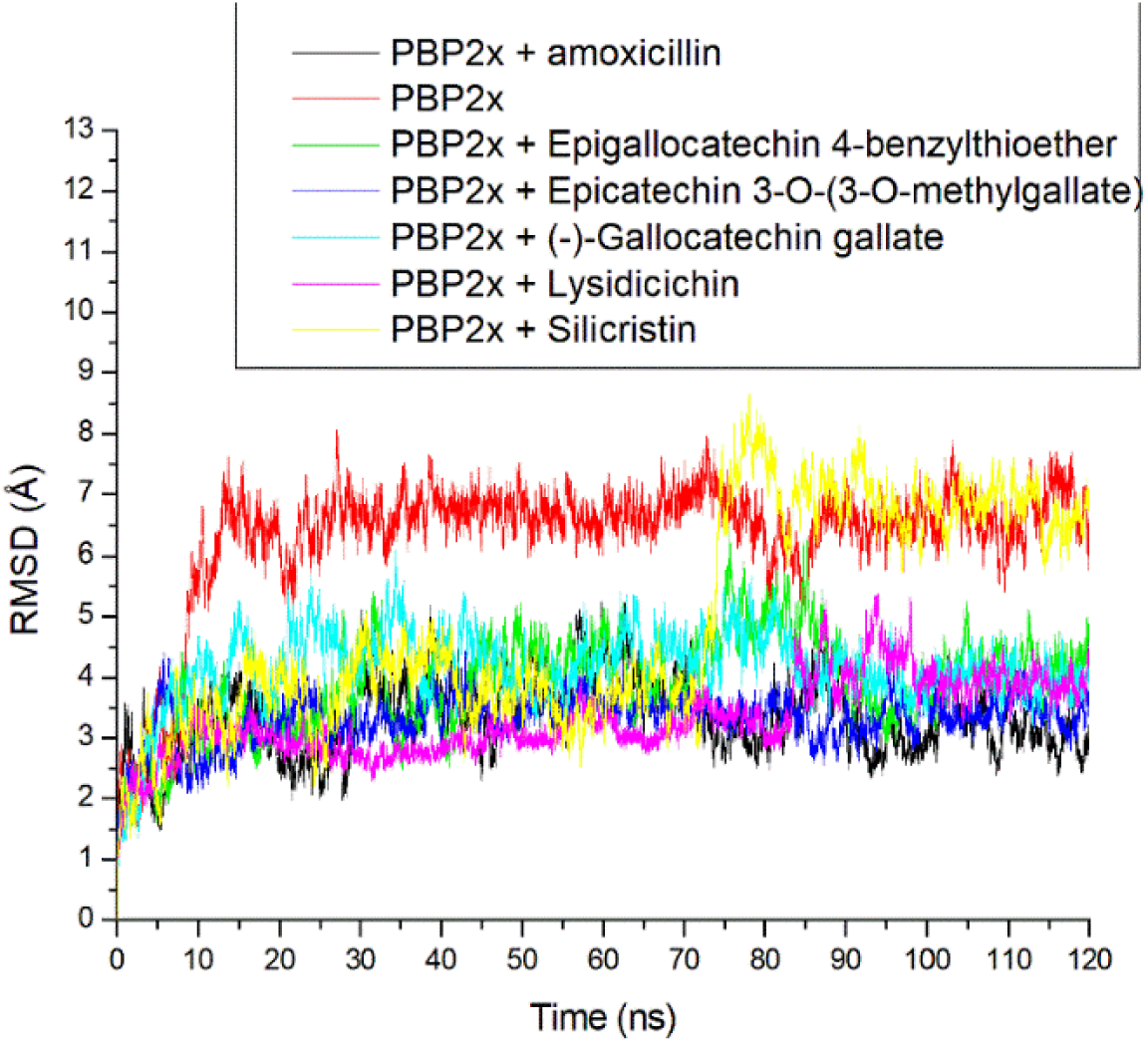
Relative root means square deviation plots of alpha-carbon, amoxicillin, and top-hit phenolics at the actv site of PBP2x of *S. pneumoniae* over a 120 ns simulation period.

**Table 1:** Physicochemical, pharmacokinetics and toxicity profile of the top-hit phenolics.

| Ligands zinc code | Popular Name | Code | MW <500 (g/mol) | HB-A ≤ 10 | HB-D ≤ 5 | Log Po/w ≤ 5 | RT-B ≤ 9 | WS | GI - A | BS | pgp | Inhibitor of CYP 450s |  |  |  |  | H | C | IM | M | CY | TC |
| --- | --- | --- | --- | --- | --- | --- | --- | --- | --- | --- | --- | --- | --- | --- | --- | --- | --- | --- | --- | --- | --- | --- |
|  |  |  |  |  |  |  |  |  |  |  |  | CYP 1A2 | CYP 2C19 | CYP 2C9 | CYP 2D6 | CYP 3A4 |  |  |  |  |  |  |
| Amoxicillin | Amoxicillin | - | 365.4 | 6 | 4 | -0.39 | 5 | VS | L | 0.55 | N | N | N | N | N | N | I | I | I | I | I | 6 |
| ZINC03793048 | Silicristin | - | 482.44 | 10 | 6 | 1.42 | 4 | S | L | 0.55 | N | N | N | N | N | Y | I | I | A | I | I | 4 |
| ZINC38337967 | Epigallocatechin 4-benzylthioether | ECBE | 428.46 | 7 | 6 | 2.22 | 4 | MS | L | 0.55 | N | N | N | N | N | Y | I | I | I | I | I | 5 |
| ZINC14642669 | Lysidicichin | - | 470.43 | 10 | 5 | 1.94 | 6 | MS | L | 0.55 | N | N | N | N | N | Y | I | I | A | I | I | 4 |
| ZINC14642643 | Epicatechin 3-O-(3-O-methylgallate) | ECMG | 456.4 | 10 | 6 | 1.77 | 5 | S | L | 0.55 | N | N | N | N | N | Y | I | I | A | I | I | 4 |
| ZINC03870413 | (-)- Gallocatechin gallate | GCG | 458.37 | 10 | 8 | 1.01 | 4 | S | L | 0.17 | N | N | N | N | N | N | I | I | I | I | I | 4 |

Keywords: Molecular weight = MW; Hydrogen bond acceptor = HB-A; Active = A; Permeability glycoprotein substrate = Pgp; Hydrogen bond donor = HB-D; Partition
coefficient = Log Po/w; Very soluble = VS; Rotatable bond = RT-B; Water solubility = WS; Cytotoxicity = CY; Gastrointestinal absorption = GI- A; Cytochrome = CYP;
Moderately soluble = MS; Soluble = S; No = N; Toxicity class = TC; Yes = Y; Low = L; Inactive = I; Bioavailability score = BS; Hepatotoxicity = H; Carcinogenicity = C;
Immunotoxicity = IM; Mutagenicity = M.

### 2.2 Energy components refinement of the top-hit phenolics against the actv site of PBP2x

The molecular docking study has been shown to be a poor predictor of binding affinity and the identification of active inhibitors due to its scoring functions inability to account for the dynamics of the atoms of the ligand and the protein involved [21]. Thus, in this study, energy refinement of the docking score was done using the MMGBSA approach [26]. Using this approach, the ΔGbind, which gives the sum of all the intermolecular interactions that are present between a ligand and a target during a simulation, was calculated. Several energy components, such as the ΔE_elec_, ΔE_gas_, ΔE_vdw_ and ΔG_sol_ contributed to the total ΔGbind of a ligand to a receptor. In this study, the ΔG_sol_, which estimates the the electrostatic solvation of the solute [27], had the highest contribution to the total ΔGbind observed for all the investigated complexes (Table 2). Three of the top-hit phenolics [ECMG (−42.18 kcal/mol), silicristin (−38.41 kcal/mol), and lysidicichin (−29.85 kcal/mol)] had higher ΔGbind than amoxicillin, with ECMG having the highest negative value (Table 2). This observation, is not only consistent with previous studies on the antibacterial properties of phenolics [17], but indicative of the enhanced potential of the three phenolics as modulators of PPBP2x, with ECMG being the most promising compound. ECBE (−19.72 kcal/mol) had ΔGbind scores that compared with amoxicillin (−19.23 kcal/mol), while the value observed for GCG (−14.93 kcal/mol) was significantly lower relative to amoxicillin (Table 2). This observation is suggestive of the lesser ability of ECBE and most especially GCG to modulate PBP2x of *S. pneumoniae*, thus contradicting the observed docking scores, where the two compounds had higher scores than amoxicillin. This finding substantiates the report of Cerón-Carrasco [21] that virtual screening required further energy refinement in the selection of active molecules. Overall, the findings from the thermodynamic ΔGbind of this study demonstrated the benefit of ECMG, silicristin, and lysidicichin over amoxicillin in the management/treatment of infections caused by multidrug resistant *S. pneumoniae*.

**Table 2.** Thermodynamic Energy (kcal/mol) of the top-five phenolics against the actv site of PBP2x of *S. pneumoniae*.

| Systems | $\Delta E_{vdW}$ | $\Delta E_{elec}$ | $\Delta G_{gas}$ | $\Delta G_{solv}$ | $\Delta G_{bind}$ |
| --- | --- | --- | --- | --- | --- |
| Amoxicillin | $-29.65 \pm 4.89$ | $-290.87 \pm 33.31$ | $-320.52 \pm 34.44$ | $301.29 \pm 27.75$ | $-19.23 \pm 7.96$ |
| Silicristin | $-46.96 \pm 4.46$ | $-41.91 \pm 9.87$ | $-88.90 \pm 9.57$ | $50.48 \pm 7.77$ | $-38.41 \pm 6.07$ |
| ECBE | $-23.59 \pm 8.08$ | $-22.28 \pm 14.23$ | $-45.88 \pm 18.96$ | $26.15 \pm 12.22$ | $-19.72 \pm 8.15$ |
| Lysidicichin | $-37.47 \pm 7.64$ | $-32.83 \pm 10.17$ | $-70.30 \pm 13.40$ | $40.45 \pm 7.24$ | $-29.85 \pm 8.56$ |
| ECMG | $-49.42 \pm 3.96$ | $-45.24 \pm 8.76$ | $-94.66 \pm 8.28$ | $52.48 \pm 5.19$ | $-42.18 \pm 4.53$ |
| GCG | $-16.35 \pm 13.69$ | $-27.02 \pm 22.28$ | $-43.37 \pm 29.16$ | $28.43 \pm 19.81$ | $-14.93 \pm 10.27$ |

Keyword: $\Delta E_{vdW}$ means van der Waals energy, $\Delta G_{bind}$ means total binding free energy, $\Delta E_{gas}$ means gas phase free energy, $\Delta E_{elec}$ means electrostatic energy, while $\Delta G_{solv}$ means solvation free energy

### 2.3 Thermodynamic information of PBP2x following binding at the actv site

To understand the structural stability of PBP2x and its complexes after ligand binding, the RMSD was assessed. The RMSD is a function of the time-dependent deviation of a complex structure from its apo structure, with lower RMSD values indicating better complex stability [28]. In this study, the RMSD seems to equilibrate before 10 ns for all the investigated systems, while each system subsequently took a more diverse route, which appears to converge at 85 ns of simulation, thus impacting the different average RMSD values observed for each system (Figure 2). The bound systems with average values ranging between 3.31 and 4.90 Å, generally had a reduced RMSD value compared to the apo-PBP2x (6.32 Å) with lysidicichin and ECMG having the lowest values, which compared well with amoxicillin (Table 3). The reduced RMSD value of PBP2x following binding of ligands generally denotes enhanced thermodynamic stability of the complexes [29]. This observation agrees with the report of Sainsbury *et al*. [30], where experimental beta-lactam binding of PBP3 increases the thermodynamic stability of PBP3 through local conformational changes at the actv site. The fact that lysidicichin and ECMG had RMSD values that compared well with amoxicillin following binding of PBP2x indicates their advantage as inhibitors of PBP2x over the other top-hit phenolics. This observation partially agrees with the ΔGbind findings of this study, where ECMG had the highest negative ΔGbind. However, silicristin, which also had higher ΔGbind against PBP2x, had the highest RMSD value at 4.90 Å (Table 3) and thus denotes that further structural adjustment of the compound for stability might improve potency. Overall, the RMSD finding of the study showed that ligand binding of PBP2x increases stability when compared to the apo-PBP2x and that while lysidicichin and ECMG are the most structurally compactible with PBP2x, silicristin and probably GCG might require further structural adjustment to increase their potency.

**Table 3.** Thermodynamics data of the top-five phenolics after 120 ns MD simulation at the actv site of PBP2x of *S. pneumoniae*.

| Systems | RMSD (Å) | RMSF (Å) | ROG (Å) | SASA (Å) | Number of H-bonds | Average distance of H-bonds |
| --- | --- | --- | --- | --- | --- | --- |
| Apo-PBP2x | $6.32 \pm 1.08$ | $1.76 \pm 1.02$ | $30.04 \pm 0.17$ | $27812.69 \pm 465.30$ | $328.68 \pm 1.02$ | $2.85 \pm 0.04$ |
| Amoxicillin | $3.33 \pm 0.61$ | $2.16 \pm 1.35$ | $29.37 \pm 0.50$ | $28553.79 \pm 520.35$ | $329.78 \pm 1.12$ | $2.85 \pm 0.05$ |
| Silicristin | $4.90 \pm 1.72$ | $2.62 \pm 2.18$ | $29.63 \pm 0.41$ | $28080.16 \pm 436.40$ | $333.65 \pm 1.05$ | $2.85 \pm 0.04$ |
| ECBE | $3.88 \pm 0.78$ | $1.83 \pm 1.15$ | $29.51 \pm 0.23$ | $28586.34 \pm 465.61$ | $331.34 \pm 0.99$ | $2.85 \pm 0.02$ |
| Lysidicichin | $3.31 \pm 0.61$ | $1.76 \pm 1.00$ | $29.15 \pm 0.19$ | $28065.41 \pm 565.10$ | $331.32 \pm 0.98$ | $2.85 \pm 0.04$ |
| ECMG | $3.32 \pm 0.39$ | $1.64 \pm 1.01$ | $29.61 \pm 0.22$ | $28612.65 \pm 434.72$ | $333.98 \pm 1.13$ | $2.85 \pm 0.03$ |
| GCG | $4.14 \pm 0.62$ | $1.85 \pm 2.08$ | $30.01 \pm 0.25$ | $28378.68 \pm 453.43$ | $330.02 \pm 1.08$ | $2.85 \pm 0.06$ |

The ROG estimates the degree of folding with great folding meaning better thermodynamic organization and sometimes also stability in a complex, a phenomenon related to protein (topology, size, and length) and ligand (molecular weight) parameters and their point of contact [31]. Like the RMSD, following ligand binding of PBP2x, a marginal reduction in the ROG values was observed with the bound systems relative to the apo-PBP2x (Table 3, Figure 3). The reduced ROG value following ligand binding of PBP2x could be suggestive of the increased structural compactness in the bound systems due to folding [32]. The bound system had average ROG values ranging between 29.15 and 30.01 Å with lysidicichin having the lowest value (Table 3). Lysidicichin having the lowest ROG value agrees with the RMSD findings of this study, where the compound also had the lowest RMSD value, and thus suggests the advantage of the compound as a potential PBP2x inhibitor. However, with a variance of 0.86 Å (Table 3) among the bound systems regarding the average ROG, it can be inferred that the compactness of PBP2x following ligand binding had little impact on ranking the top-hit phenolics.

**Figure 3:**
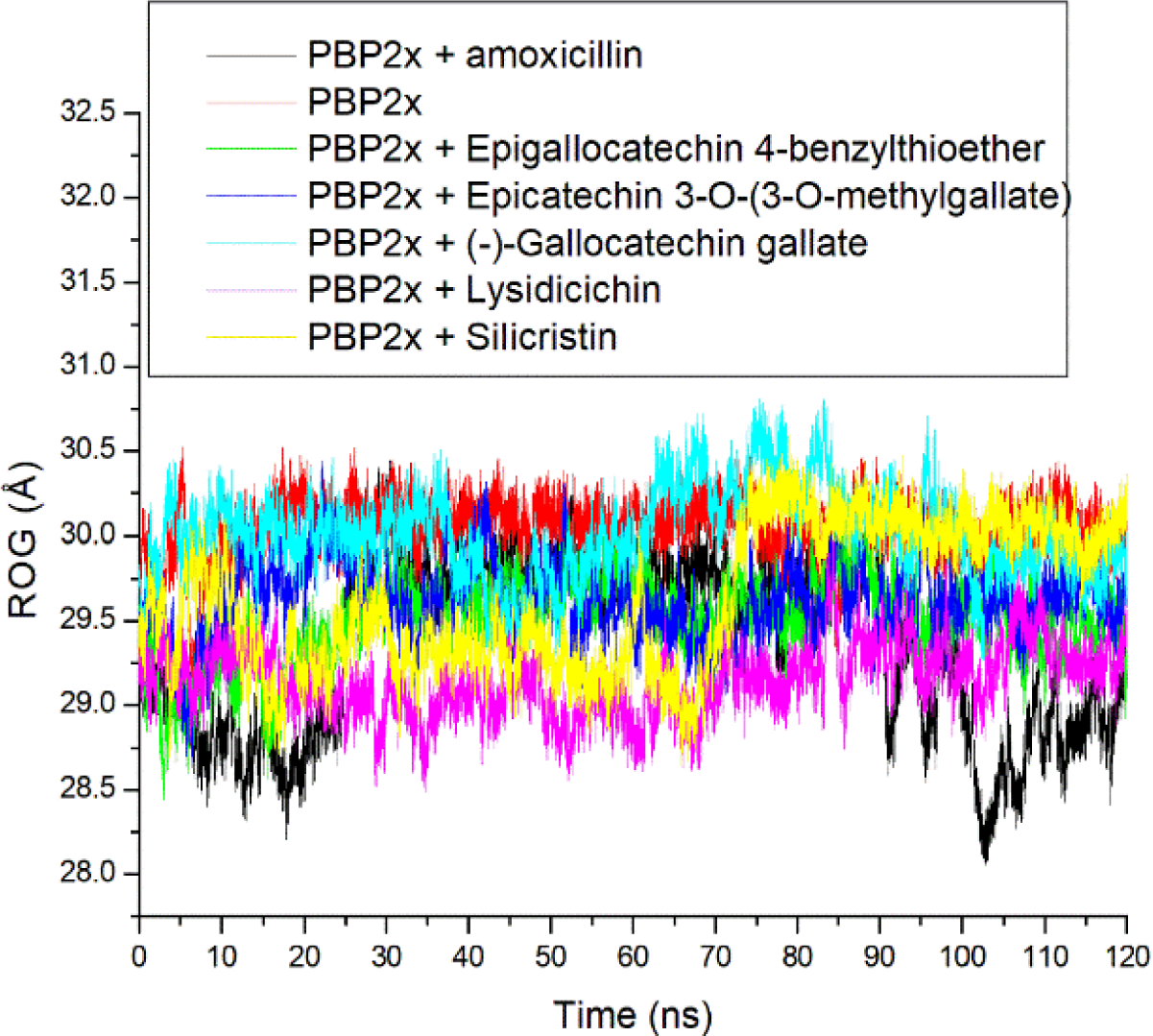
Relative radius of gyration plots of alpha−carbon, amoxicillin and top-hit phenolics at the actv site of PBP2x of *S. pneumoniae* over a 120 ns simulation period

Root mean square fluctuation evaluates the mobility of residues following ligand binding of PBP2x [33]. In this study, higher swaying in amino acid residues between 25 and 175 and a lesser fluctuation between residues 250 and 425, and 550 and 625 were noted in all the systems, with the degree of swaying varying among systems (Figure 4a), thus affecting the average RMSF. Interestingly, Ser337, which is the catalytic residue in PBP2x [10], occurred where the fluctuation was minimal (Figure 4b). Ser337 had fluctuations ranging between 0.87 Å and 1.29 Å in all the system with the apo-PBP2x having the highest fluctuation of Ser337 (Table 4). This observation suggests that ligand binding generally reduced the fluctuation of Ser337, indicating an active involvement of Ser337 in the intramolecular binding of other residues of PBP2x or the intermolecular binding of the ligands. The reduced effect of ligand binding on the Ser337 of PBP2x is in agreement with our previous report, where the gatekeeper residue (Tyr446) of PBP2a had reduced fluctuation following binding of phenolics [34]. Silicristin (0.87 Å) and ECMG (0.88 Å) with the highest negative ΔGbind against PBP2x also had the lowest fluctuation of Ser337, pinpointing the importance of reduced fluctuation of the catalytic residue in the observed ΔGbind in the study. However, except for the ECMG-PBP2x complex (1.64 Å), the fluctuation of residues in the whole protein was greatly enhanced following the top-hit phenolics and amoxicillin binding of PBP2x (Table 3). The apo-PBP2x had an RMSF of 1.76 Å, while the bound systems had average RMSF values ranging between 1.64 Å and 2.62 Å with only ECMG-PBP2x having a lesser value than the apo-PBP2x. The lesser RMSF value of ECMG-PBP2x suggests enhanced structural organization with the binding domain of PBP2x due to strong intra and intermolecular binding and could have contributed to the higher negative ΔGbind observed in ECMG-PBP2x relative to other investigated complexes in this study. Like the findings on the RMSD, the silicritin-PBP2x complex had the highest fluctuation of residues in the whole protein, suggesting the need for structural adjustment of the compound for stability. Other top-hit phenolic complexes with PBP2x, such as ECBE-PBP2x (1.83 Å), lysidicichin-PBP2x (1.76 Å), GCG-PBP2x (1.85 Å) had relatively lower RMSF values than amoxicillin-PBP2x (2.16 Å) (Table 3), suggesting the advantage of ECBE, lysidicichin, and GCG as potential inhibitors of PBP2x. Overall, the RMSF findings of this study have demonstrated the importance of reduced fluctuation of Ser337 in the observed ΔGbind noted in this study and that ECMG among the investigated ligands caused enhanced structural organization in PBP2x, which could have contributed to its higher negative ΔGbind. Silicristin, on the other hand, despite having a high ΔGbind, might require further structural adjustment to increase its potency due to its high RMSF value, and this observation is also noted by the RMSD finding of this study.

**Figure 4:**
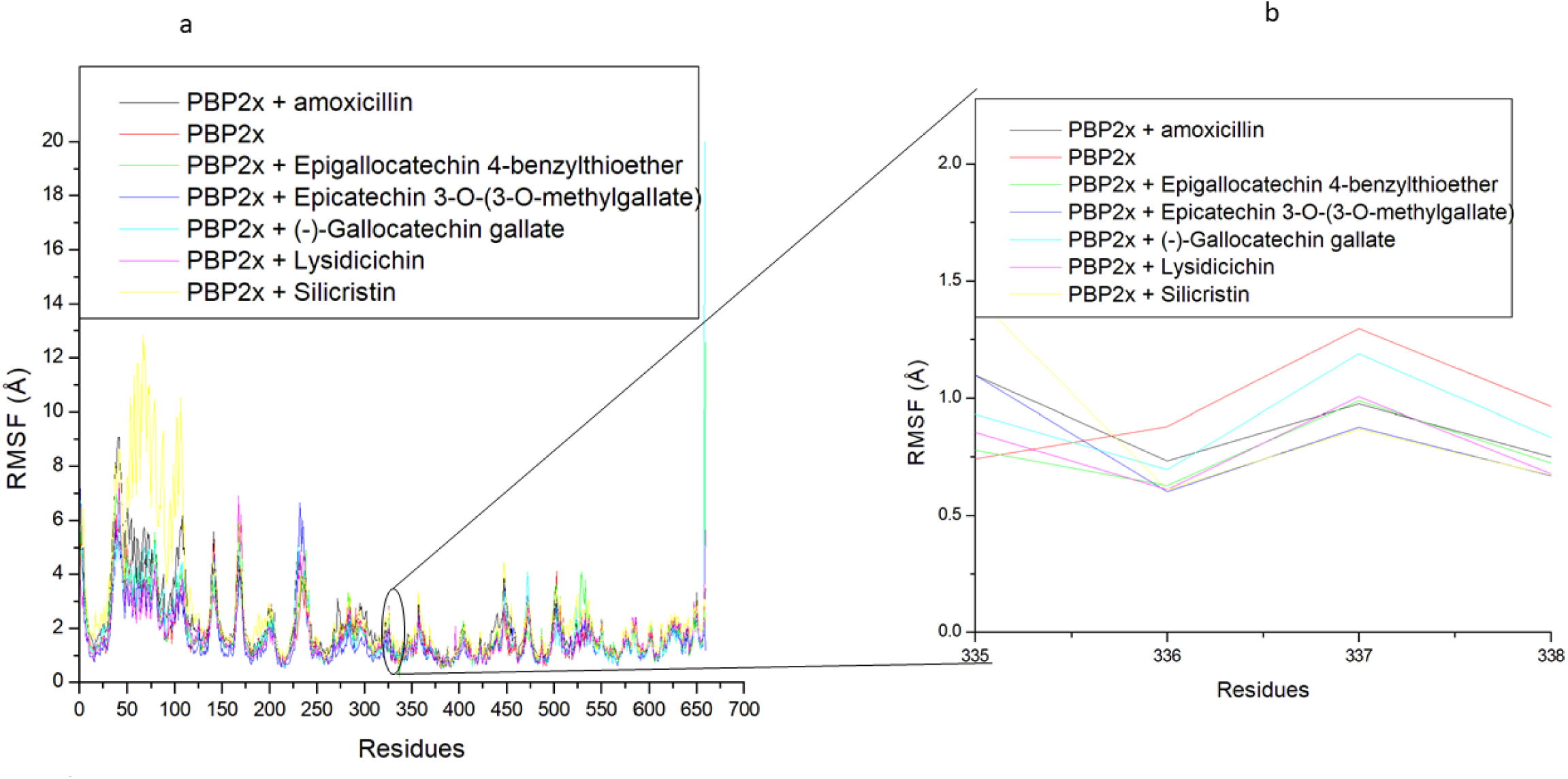
Relative root means square fluctuation plots of alpha−carbon, amoxicillin and top-hit phenolics at the actv site of PBP2x (a) and the critical residue of PBP2x (b) of *S. pneumoniae* over a 120 ns simulation period

**Table 4:** PBP2x actv site Ser337 fluctuation (Å) after 120 ns simulation of the top-hit phenolics at the actv site.

| PBP2x actv site critical residue | Top hit phenolics |  |  |  |  |  |  |
| --- | --- | --- | --- | --- | --- | --- | --- |
|  | Apo-PBP2x | Amoxicillin | Silicristin | ECBE | Lysidicichin | ECMG | GCG |
| Ser337 | 1.29 | 0.97 | 0.87 | 0.99 | 1.00 | 0.88 | 1.19 |

Like the ROG, the SASA also analyzes protein folding and surface area changes during a simulation, with higher SASA values indicating a rise in volume [35]. The SASA plot had a more stable fluctuation throughout the simulation for all the systems (Figure 5). Averagely, the bound system had slightly higher SASA values ranging between 28065.41 Å and 28612.65 Å compared to the apo-PBP2x, with lysidicichin having the lowest value (Table 3). This observation highlighted the slightly higher surface area of PBP2x following ligand binding, which is in contrast with the findings of Mousavi et al. [33], who reported that the interactions of ligands with proteins can cause protein folding. The increase in surface area following ligand binding also did not agree with the ROG findings of this study. This observation might not necessarily mean less stability of the complexes, as a smaller SASA variance of 800 Å exists between the bound system and the apo-PBP2x (Table 3). However, lysidicichin, among the investigated compounds, has the lowest SASA value, which agrees with the ROG and RMSD findings of this study and thus further corroborates the advantage of the compound as PBP2x inhibitor. Overall, with a smaller SASA variance of 547 Å among the bound systems, it can be concluded that the surface area folding of PBP2x following ligand binding had little impact on ranking the top-hit phenolics.

**Figure 5:**
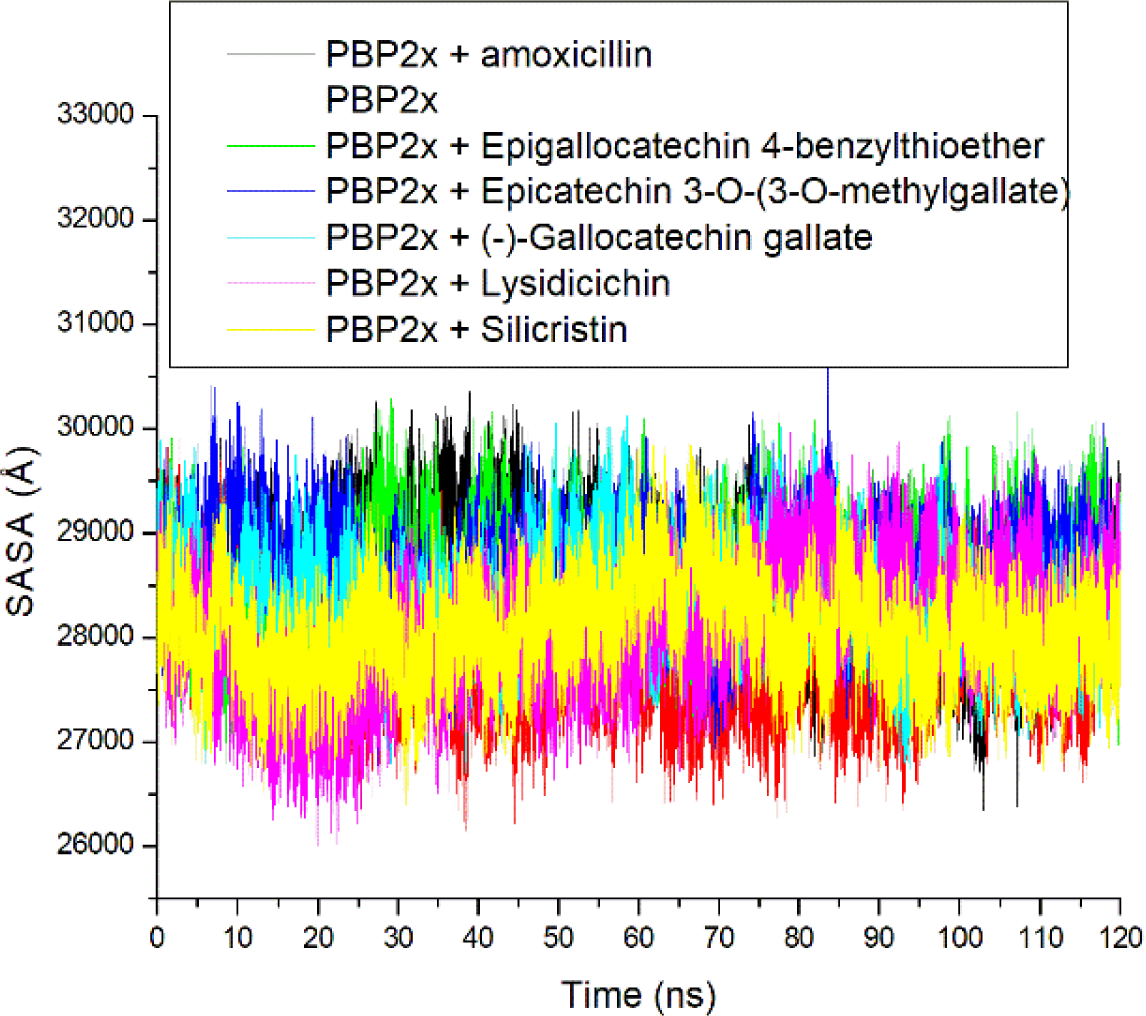
Relative solvent−accessible surface area plots of alpha−carbon, amoxicillin and top-hit phenolics at the actv site of PBP2x of *S. pneumoniae* over a 120 ns simulation period

Hydrogen bonds and distances are important for the structural stability of proteins and can hence be evaluated during simulation to understand the influence of ligand binding on protein stability [36]. A stable fluctuation in the number of intramolecular hydrogen bonds was observed with bonded and apo-PBP2x during the simulation (Figure 6a). This observation, while being consistent with other thermodynamic metrics of this study, further indicates the thermodynamic orderliness of the systems [37]. Following ligand binding of PBP2x, a higher number of hydrogen bonds in PBP2x was observed with the bound system (ranging between 329.78 and 333.98) relative to the apo-PBP2x (Table 3), with ECMG having the highest number of hydrogen bonds. The increase in the hydrogen bond of the bound system relative to the apo-PBP2x is indicative of some intermolecular hydrogen bonds contributed by ligand binding of the protein, with ECMG having the highest intermolecular contribution of hydrogen bonds during the simulation. This observation could have contributed to ECMG having the highest negative ΔGbind against PBP2x and the higher negative ΔGbind observed with silicristin, ECBE, lysidicichin, and ECMG relative to amoxicillin. The relatively similar pattern of fluctuations in the distance of hydrogen bonds of PBP2x [with an average of 2.85 Å for each of the systems (Table 3)] following the top-hit phenolics binding of PBP2x, which continues to shorten as the simulation progresses (Figure 6b) also further emphasized the orderliness in the protein following the compounds binding at the actv site, an observation that agrees with other thermodynamic metrics of this study.

**Figure 6:**
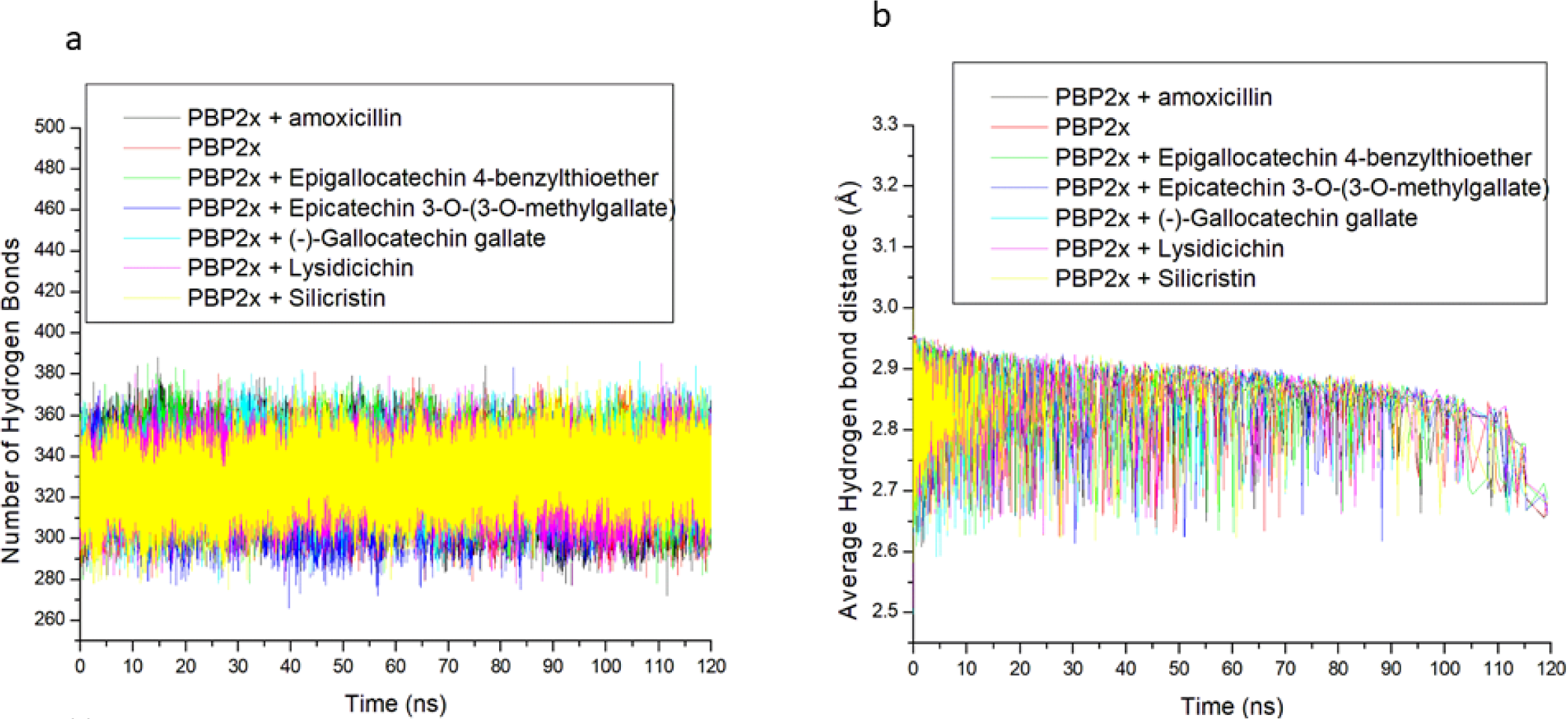
Time progression of (a) number of H-bonds, and (b) distance formed in PBP2x following ligand binding at the actv site during the simulation.

### 2.4 Molecular docking of top hit phenolics at the allo site of PBP2x of S. pneumoniae

The PASTA domain of PBP2x has been demonstrated to act as an allo site, which, when properly occupied, could facilitate entering the actv site of PBP2x and thus prompt the acylation of Ser337 of the actv site [12]. To investigate the ability of the top-ranking phenolics binding the allo site of PBP2x, molecular docking was carried out to assess the fitness of the phenolics at the site. The top-hit phenolics had higher negative docking scores at the allo site, ranging between −9.1 and −9.9 kcal/mol, than amoxicillin (−8.8 kcal/mol), with silicristin having the highest negative docking scores (Table 5). This observation might be indicative of the better fitness of the top-hit phenolics relative to amoxicillin at the allo site of PBP2x, with silicristin showing the better fitness. The docking scores observed at the allo site are comparable with those observed at the actv site, which also range between −9.2 and −9.9 kcal/mol, with silicristin having the highest negative docking scores at both sites. However, as molecular docking approach only serves as a screening process for obtaining the pose, orientation, and interactions of a ligand at the binding pocket of a protein, MMGBSA and molecular dynamic studies were further carried out to assess the ΔGbind, thermodynamic stability, and fluctuation of the actv site Ser337 following binding of the PBP2x allo site by the top-hit phenolics.

**Table 5:** Molecular docking, thermodynamics ΔGbind, whole RMSF and RMSF values of Ser337 following binding of top hit phenolics at the allo site of PBP2x of *S. pneumoniae*.

| Systems | Docking score (kcal/mol) | $\Delta G_{bind}$ (kcal/mol) | RMSF (Å) | RMSF of Ser337 (Å) |
| --- | --- | --- | --- | --- |
| Apo-PBP2x |  |  | 13.05 ± 0.79 | 11.44 ± 0.12 |
| Amoxicillin | -8.8 | -33.75 ± 9.87 | 10.56 ± 0.77 | 14.97 ± 0.30 |
| Silicristin | -9.9 | -50.54 ± 4.88 | 7.86 ± 0.79 | 18.75 ± 0.22 |
| ECBE | -9.3 | -45.82 ± 9.74 | 10.78 ± 0.64 | 16.59 ± 0.23 |
| Lysidicichin | -9.2 | -33.53 ± 5.84 | 8.43 ± 0.79 | 15.48 ± 0.24 |
| ECMG | -9.1 | -35.06 ± 6.12 | 15.79 ± 0.74 | 14.03 ± 0.12 |
| GCG | -9.4 | -44.21 ± 5.74 | 9.61 ± 0.64 | 16.60 ± 0.23 |

### 2.5 Thermodynamic ΔGbind, stability and compactness following simulation of the top-ranked phenolics at the allo site of PBP2x of S. pneumoniae

Following energy refinement of the top-hit phenolics at the allo site of PBP2x, silicristin (−50.54 kcal/mol), ECBE (−45 kcal/mol), ECMG (−35.06 kcal/mol), and GCG (−44.21 kcal/mol) had higher negative ΔGbind at the allo site of PBP2x than amoxicillin (−33.75 kcal/mol), with silicristin having the highest negative score (Table 5, Table S2). This observation is suggestive of the better affinity of the top-hit phenolics for the allo site of PBP2x, with silicristin being the most promising as a probable inhibitor at the site, and interestingly, this finding agrees with the docking findings of this study. The observed high affinity of silicristin at both the actv and allo sites of PBP2x points to the probable broad-spectrum activity of the compound as an inhibitor that can treat or act in synergy with conventional antibiotics in the treatment of infections caused by *S. pneumoniae*. This observation is consistent with previous studies where phenolics show good synergistic properties with conventional antibiotics [14 19]. ECBE and GCG also had good ΔGbinds at the allo site of PBP2x, indicating their probable capability to act in synergy with conventional beta-lactams (Table 5). ECMG and lysidicichin, on the other hand, had moderately higher negative ΔGbinds than amoxicillin (Table 5). Analysis of the thermodynamic stability suggests stable trajectories of the top-hit phenolics at the allo site of PBP2x with RMSD values that are less than 3 Å limit [38] (ranging between 1.74 and 2.19 Å) and negligible ROG and SASA variances of 0.23 Å and 1042 Å respectively, between the bonded systems and the apo-PBP2x (Figure S2, Table S3). These observations are revealing of the thermodynamic compatibility of the top-hit phenolics at the allo site of PBP2x. While two different experimental structures of PBP2x were used for the actv and allo site evaluations, the low RMSD values and smaller ROG variance following binding at the allo site relative to the actv site could be pinpointing the higher thermodynamic stability of the top-hit phenolics at the allo site.

### 2.6 Thermodynamic fluctuation of the actv site Ser337 following simulation of the top-ranked phenolics at the allo site of PBP2x of S. pneumoniae

The RMSF fluctuation of PBP2x following the top-hit phenolics and amoxicillin binding of PBP2x ranges between 9.61 and 15.79 Å with silicristin having the lowest value (Table 5, Figure 7a). Unlike the actv site, this observation implies a significantly higher fluctuation of the residues of PBP2x following the top-hit phenolics and amoxicillin binding at the allo site. However, it is worth acknowledging that the different experimental crystal structures of PBP2x used in this study for the actv and allo sites might have played a major role in the differences observed. Substantiating this observation is the fact that except for ECMG (15 Å), the top-hit phenolics and amoxicillin binding of PBP2x all resulted in a reduced fluctuation of residues relative to the apo-PBP2x (Table 5, Figure 7a). This observation implies that the residues were able to form stable intra- and intermolecular bonds [36] during the MD simulation of the investigated compounds at the allo site of PBP2x. Silicristin having the lowest RMSF value at 7.86 Å (Table 5) suggests the advantage of the compound as a probable inhibitor of the site, an observation that is consistent with the thermodynamic ΔGbind findings of this study. However, it is surprising that while silicristin had the lowest RMSF value in the whole protein, the compound instigated the highest fluctuation of Ser337 (Table 5, Figure 7b), a critical residue at the actv site PBP2x [10]. The highest fluctuation of Ser337, in silicristin-PBP2x complex is coherent with the highest negative ΔGbind of silicristin (−50.54 kcal/mol) at the allo site of PBP2x. Similarly, ECBE (−45.82 kcal/mol) and GCG (−44.21 kcal/mol), with the second and third highest negative ΔGbind at the allo site, had higher fluctuations of Ser337 than other investigated compounds at 16.59 Å and 16.60 Å respectively (Table 5, Figure 7b). While other investigated compounds with moderately high negative ΔGbind at the allo site also have a higher fluctuation of Ser337 than apo-PBP2x (Table 5, Figure 7b). All these observations are suggestive of the high instability of Ser337 of the actv site following ligand binding of the allo site, with the ΔGbind of the ligands at the allo site partially correlating the fluctuation of the critical amino acid. Another conclusion from this is that there is less involvement of the critical amino acid in the intramolecular binding of other important residues at the actv site, which might mean less obstruction and thus encourage the entry of beta-lactam. This observation is consistent with the earlier report of Bernardo-Garca et al. [12], who demonstrated that the complexation of a pentapeptide strain at the PASTA domains, prompts another pentapeptide strand in the same nascent peptidoglycan strand to enter the actv site of PBP2x. Another study by Mahasenan et al. [39] also showed experimentally that the displacement of adjacent residues that can enter intramolecular interactions with Tyr446 (the critical amino acid in PBP2a) after allosteric binding of PBP2a with cefepime, which has also been demonstrated computationally by Aribisala and Sabiu [34], that appropriate occupancy of the allo site of PBP2a by phenolics caused high fluctuation of Tyr446. Thus, the high fluctuation of Ser337 following ligand binding to the allo site of PBP2x is consistent with previous report. This can be further substantiated when the fluctuation of the critical amino acid is compared to the fluctuations when the top-hit phenolic and amoxicillin bind at the actv site. Binding of the top-hit phenolics at the actv site caused significant lesser fluctuation of Ser337 (between 0.87 Å and 1.19 Å) relative to the apo-PBP2x (1.27 Å) (Table 4, Figure 4b) while binding at the allo site caused significant higher fluctuation (between 14.03 and 18.75 Å) relative to the apo-PBP2x (11.44 Å) (Table 5, Figure 7b). This observation is revealing of the inter- or intramolecular involvement of Ser337 following ligand binding at the actv site, while binding at the allo site, on the other hand, causes lesser involvement of the residue in the intramolecular binding or obstruction of the actv site.

**Figure 7:**
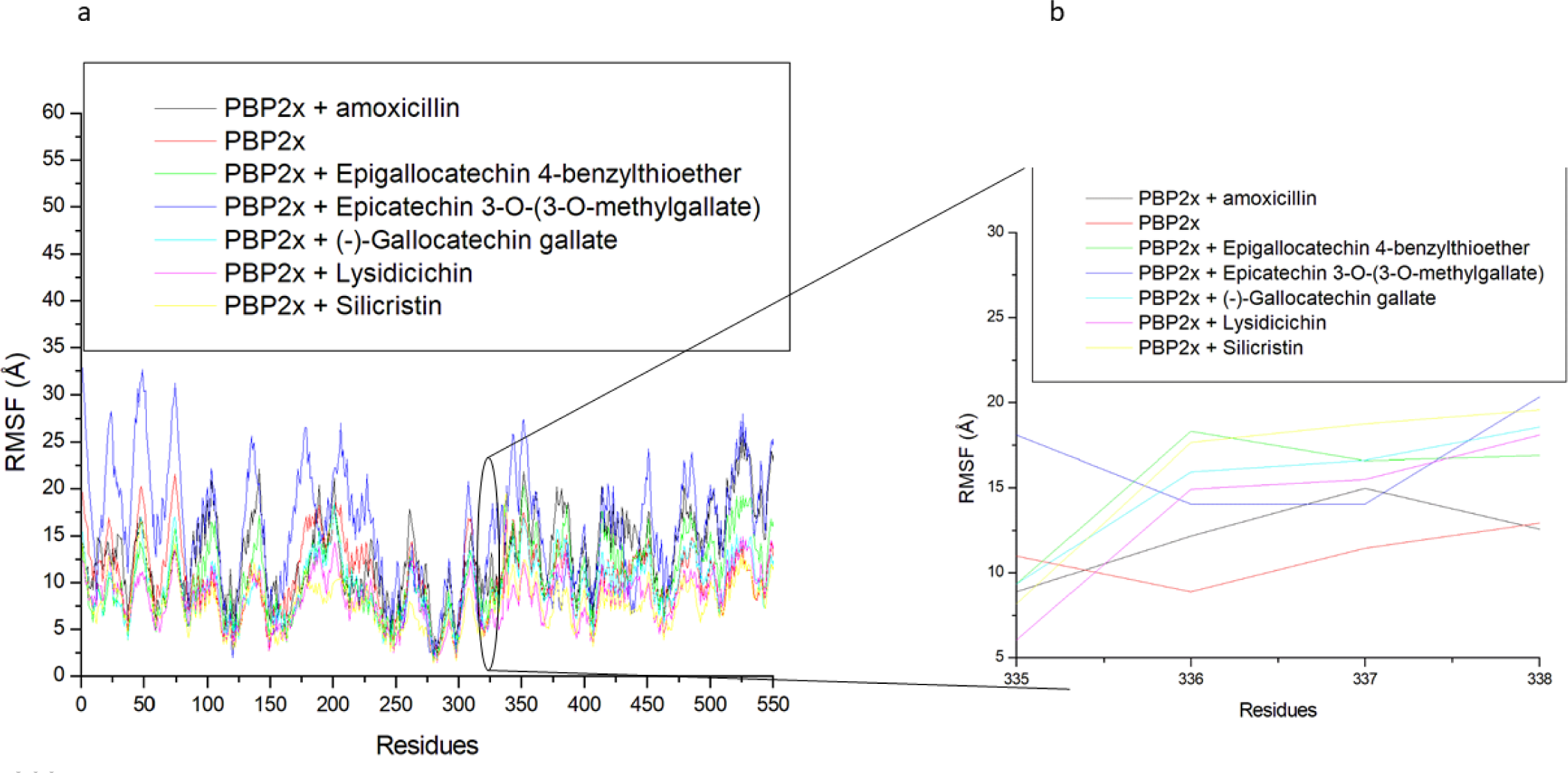
Relative root means square fluctuation plots of alpha−carbon, amoxicillin, and top-hit phenolics at the allo site of PBP2x (a) and the critical residue at the actv site of PBP2x following binding at the allo site (b) of *S. pneumoniae* over a 120 ns simulation period.

### 2.7 Interactions plots analysis of the top-hit phenolics at the actv and allo site of PBP2x of S. pneumoniae after 120 ns MD simulation

To understand how the nature of interactions impacted the ΔGbind and stability of the top-hit phenolics against PBP2x, the plots of interactions for each system after the 120 ns simulation were analyzed. The binding of drugs to biological targets has been demonstrated to rely on specific non-covalent interactions between partner molecules, including electrostatic interactions such as hydrogen bonds, van der Waals interactions, and hydrophobic effects [40]. In this study, the number, nature, and lengths of interactions between partner molecules (ligands and PBP2x) were noted to have an impact on the ΔGbind and stability observed in this study. The plots of interactions of the reference standards and phenolics with the highest ΔGbind at both the actv and allo sites of PBP2x are presented in Figures 8–11, while those of the other top-hit phenolics are shown in Figures S3 and S4 for the actv and allo sites, respectively. At the actv site, except for silicristin, epicatechin 3-O-(3-O-methylgallate), which had the highest negative ΔGbind against PBP2x interacted with the highest number of interactions at 15, three of which are hydrogen bond contacts with the lowest average distance at 2.84 Å with residues Glu295, Asn294, and Asp472 (Table 6, Figure 8). The high number of interactions and the significantly lower length of the hydrogen bond contacts could have drastically contributed to the observed high negative ΔGbind of ECMG against PBP2x. Silicristin, which has the second highest negative ΔGbind against PBP2x, had the highest number of interactions at 16, five of which are hydrogen bond contacts with an average length of 3.16 Å with residues Ser465, Tyr503, Glu295, Asn294 and Ser254 (Table 6, Figure S3). Compared to ECMG-PBP2x, even though silicristin-PBP2x had the highest number of interactions, including hydrogen bond contacts, the observed higher average distance of hydrogen bond contact in the silicristin-PBP2x complex could have contributed to its lower negative ΔGbind and stability relative to ECMG-PBP2x complex. The importance of hydrogen bond length observed in this study is consistent with the findings of Du et al. [36], who showed the relevance of reduced interaction length in the higher pull between two intramolecular or intermolecular molecules. Reduced bond length will facilitate the atoms of two molecules being held together more tightly, resulting in greater affinity and stability [36]. Similarly, lysidicichin, with a moderately higher negative ΔGbind relative to amoxicillin (Table 6, Figure 9), and other top hit phenolics had 10 interactions, 4 of which were hydrogen bond contacts with an average distance of 2.98 Å with Ser504 (2), Gln499, and Thr443 (Table 6, Figure S3). The higher number and reduced average distance of hydrogen bond contacts of lysidicichin against PBP2x could have impacted the higher negative ΔGbind relative to amoxicillin-PBP2x, ECBE-PBP2x, and GCG-PBP2x, thus further highlighting the importance of the number and bond length in ΔGbind determination. ECBE and GCG against PBP2x only had 3 and 4 total interactions with PBP2x with no hydrogen bond contacts (Table 6, Figure S3). This observation could have immensely contributed to the significantly lower ΔGbind observed with the ECBE-PBP2x and GCG-PBP2x relative to other studied complexes in this study. At the allo site, a considerable similar number of interactions, ranging from 13 to 16 total interactions, were observed between the investigated compounds and the allo site of PBP2x, with GCG having the highest interaction (Table 6). Silicristin (Figure 10), ECBE (Figure S4), and amoxicillin (Figure 11) had the same number of interactions at 15. The high number of interactions in the silicristin, GCG, and ECBE complex with PBP2x could have impacted the higher negative ΔGbind of the compounds at the allo site of PBP2x. Despite having a similar number of total interactions with silicristin and ECBE, amoxicillin has a considerably lower negative ΔGbind, which could be partly attributable to the reduced number of hydrogen bond contacts in the amoxicillin-PBP2x complex (Figure 11) relative to silicristin (Figure 10) and ECBE (Table 6, Figure S4). The catalytic residues (such as Ser337, Tyr568, Pro335, Ser396, Trp374 and Ser395 at the actv site and Arg426, Gly422, Phe421, Asn417, and Val423 at the allo site) present in the molecular docking interactions of the investigated compounds with PBP2x were not observed in the interaction plots of both sites after the 120 ns MD simulation. Similar observation was noted by Al-Karmalawy et al. [41], following simulation of alacepril with hACE2 and might have been due to the dynamic nature of simulation employed. Nevertheless, the lesser fluctuation of Ser337 (the catalytic residue in PBP2x) compared to the RMSF of the whole protein means that the top-ranked phenolics and amoxicillin during the simulation reside most at the actv region of the protein, as lesser fluctuation is tantamount to strong intermolecular bonding [42].

**Figure 8:**
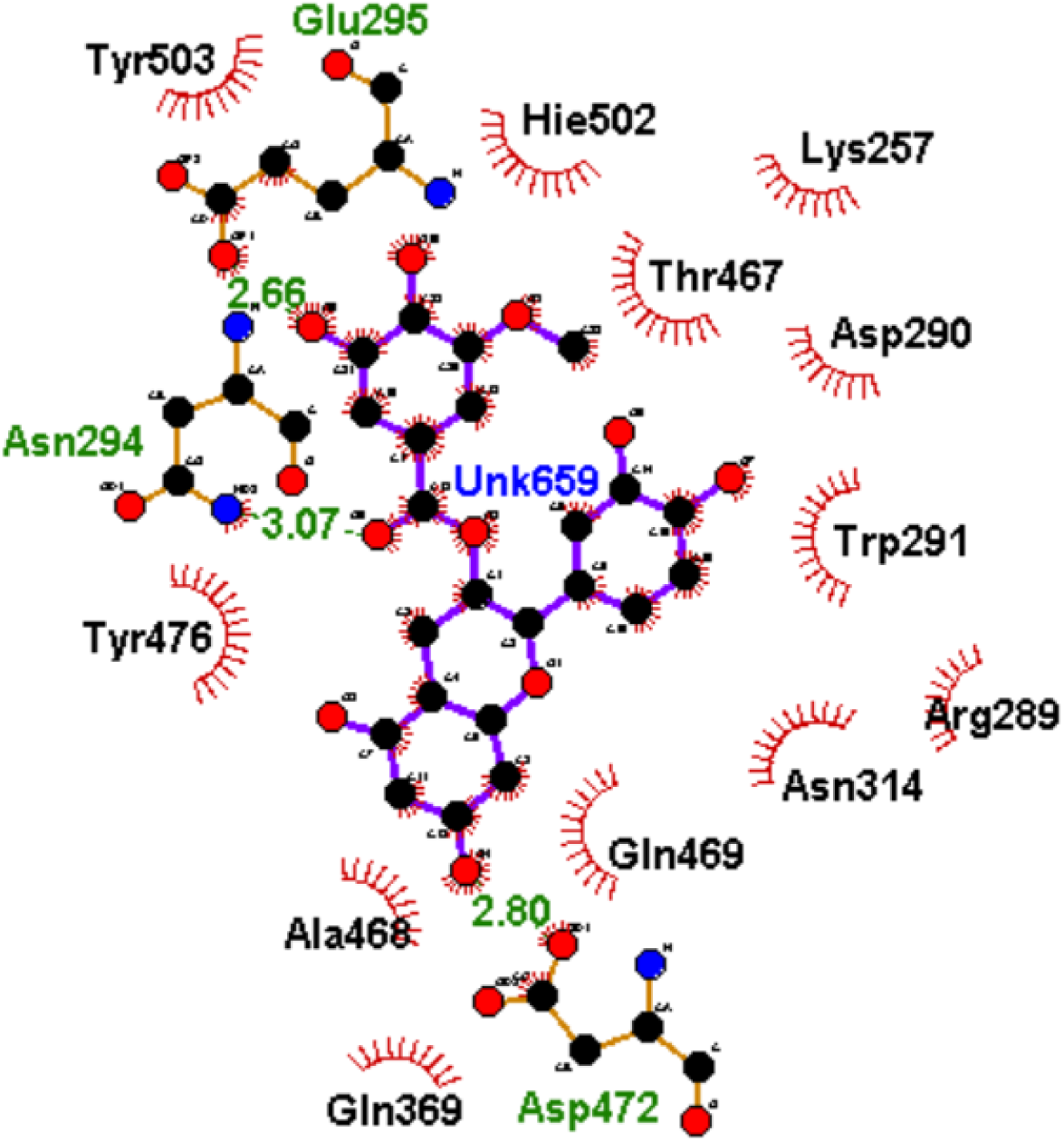
2D plot of interactions of ECMG at the actv site of PBP2x

**Figure 9:**
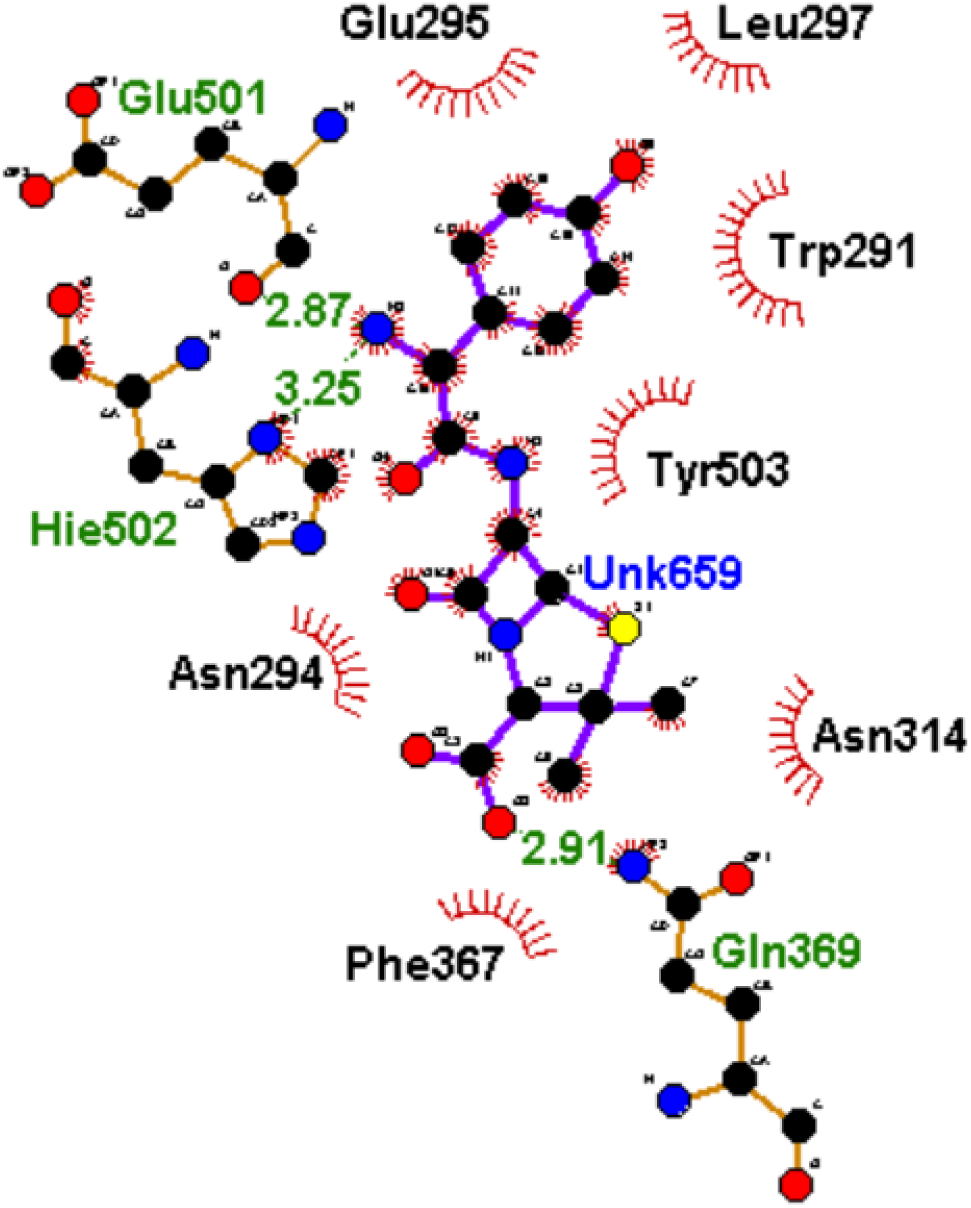
2D plot of interactions of amoxicillin at the actv site of PBP2x

**Figure 10:**
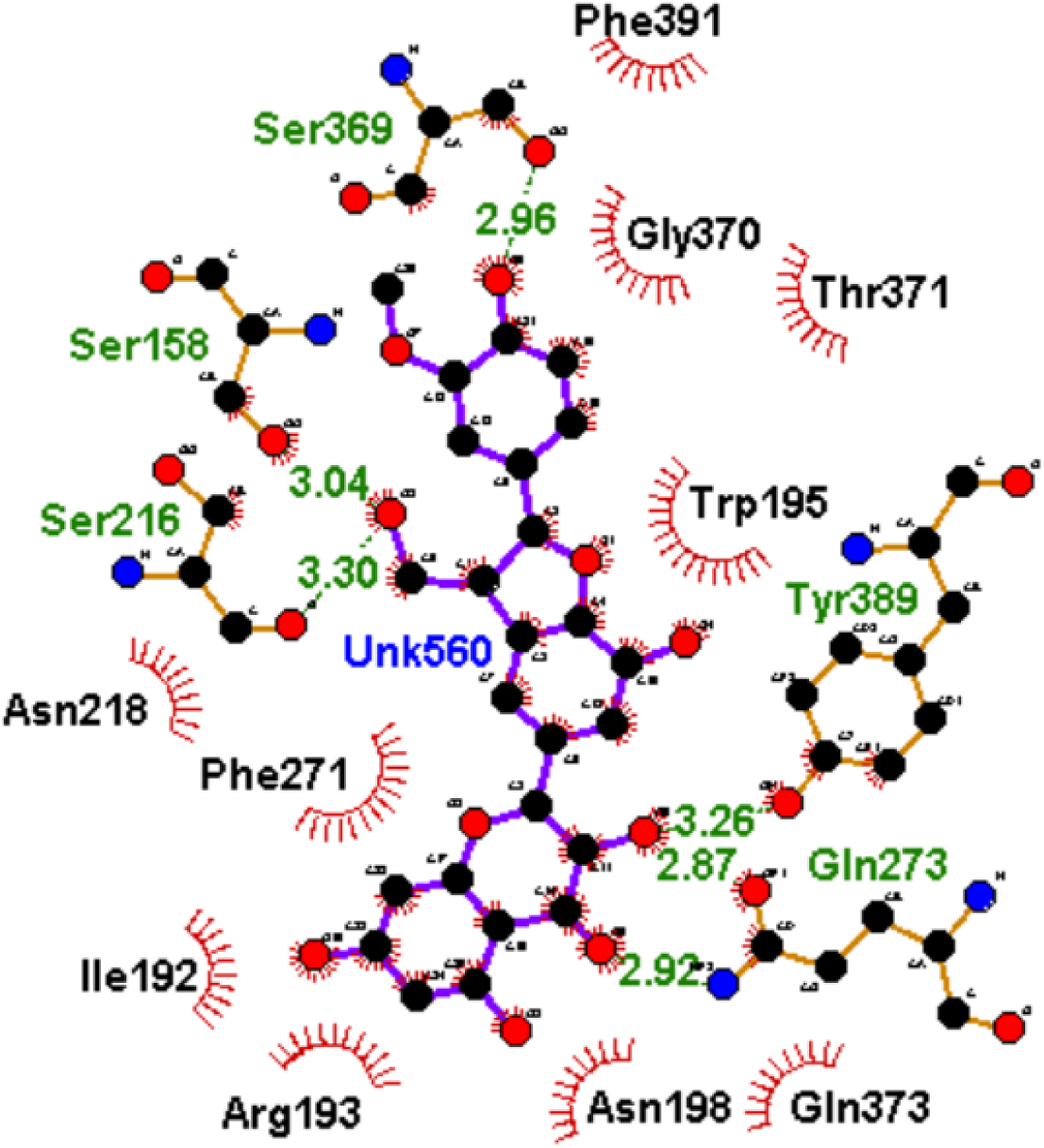
2D plot of interactions of silicristin at the allo site of PBP2x

**Figure 11:**
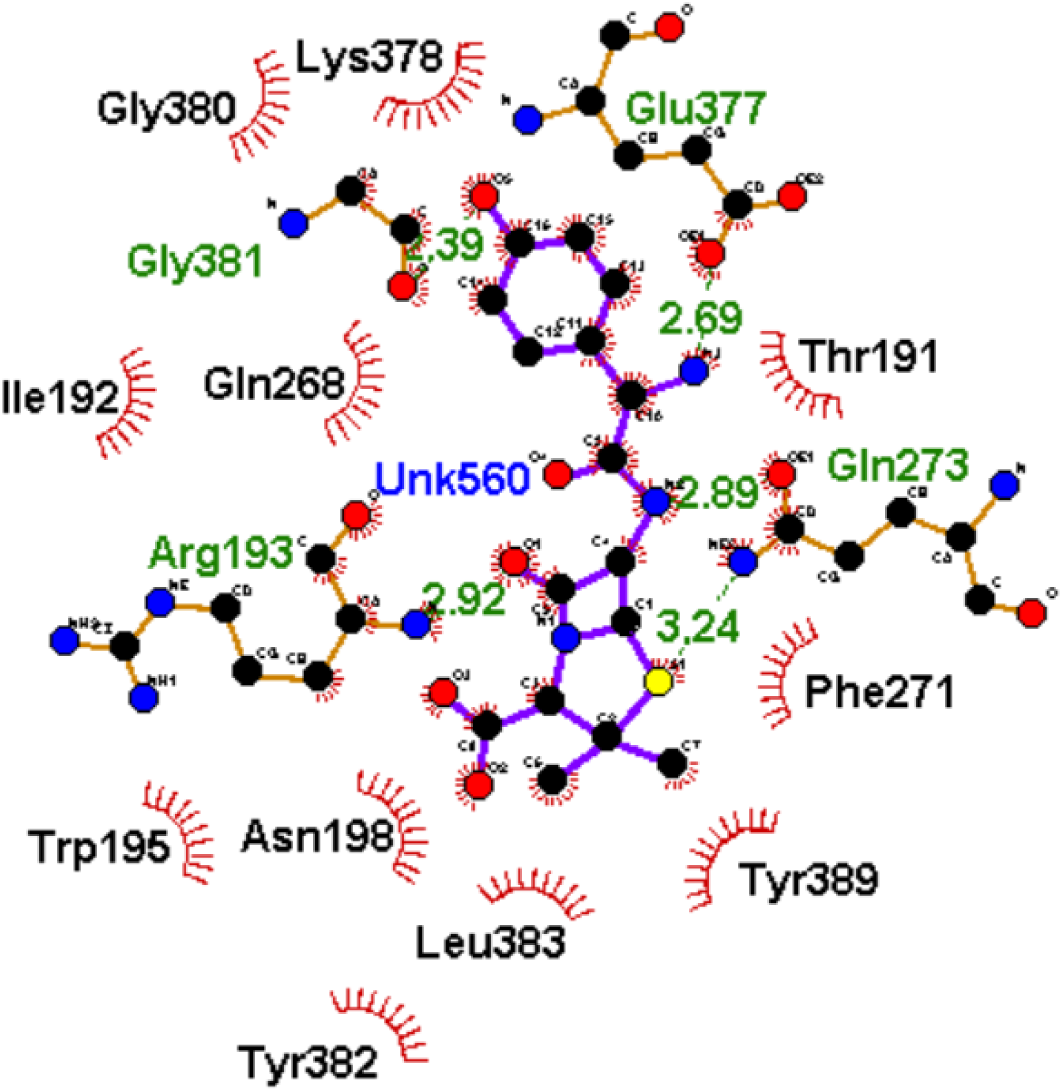
2D plot of interactions of amoxicillin at the allo site of PBP2x

**Table 6:** Interactions plots analysis of the top-hit phenolics at the actv and allo site of PBP2x of *S. pneumoniae* after 120 ns MD simulation.

| Top-ranked phenolics and standard | Total number of interactions | Number of hydrogen bonds, average distance and interaction residues |
| --- | --- | --- |
| Actv site |  |  |
| Amoxicillin | 10 | 3 (3.01 Å) [Glu501, Hie502, Asp472] |
| Silicristin | 16 | 5 (3.16 Å) [Ser465, Tyr503, Glu295, Asn294, Ser254] |
| ECBE | 3 | None |
| Lysidicichin | 10 | 4 (2.98 Å) [Ser504 (2), Gln499, Thr443] |
| ECMG | 15 | 3 (2.84 Å) [Glu295, Asn294, Asp472] |
| GCG | 4 | None |
| Allo site |  |  |
| Amoxicillin | 15 | 5 (2.82 Å) [Glu377, Gly381, Arg193, Gln273] |
| Silicristin | 15 | 6 (3.05 Å) [Gln273 (2), Tyr389, Ser216, Ser158, Ser369] |
| ECBE | 15 | 8 (2.94 Å) [Asn198, Tyr382, Ser216, Gln373, Lys161, Gln273, Phe271, Asn218] |
| Lysidicichin | 13 | 6 (2.89 Å) [Asp189, Thr191 (2), Asn262, Val264, Asp376] |
| ECMG | 13 | 6 (2.94 Å) [Asn218, Asp194 (2), Tyr381, Gly274, Glu155] |
| GCG | 16 | 3 (2.85 Å) [Leu383, Phe271, Ser158] |

### 2.8 Molecular orbital properties of best performing phenolics at the actv and allo sites of PBP2x

Understanding the HOMO and LUMO characteristics of a compound could provide crucial chemical insights relevant to the actual use of the compound as a medicinal molecule [43, 44]. The energy gap (LUMO-HOMO) of a compound provides information about its stability; A larger energy gap means high stability, which can prevent ligand binding to a protein [43, 44]. In this study, silicristin (4.0510 eV) and ECMG (4.5388 eV) had lower energy gaps than amoxicillin (5.3234 eV) (Table 7). This observation points to the high instability of silicristin and ECMG relative to amoxicillin. The higher softness and lower hardness of silicristin and ECMG relative to amoxicillin (Table 7), might suggest better reactivity of silicristin and ECMG compounds compared to amoxicillin. According to Pearson [45], higher chemical softness and lower chemical hardness signify higher compound reactivity. The high instability and better reactivity of silicristin and ECMG relative to amoxicillin could have positively impacted their better interactions at the actv and allo sites of PBP2x. Furthermore, the ability of an atom or a functional group of a compound to attract electrons could be assessed by evaluating the electronegativity and chemical potential. In this study, the higher electronegativity and higher negative chemical potential of silicristin and ECMG relative to amoxicillin (Table 7) means better capability of the atoms and functional groups of silicristin and ECMG to attracts electron at the binding pockets of PBP2x and hence their better affinity for PBP2x than amoxicillin in this study. This observation is also further supported by the higher electrophilicity index of silicristin and ECMG compared to amoxicillin (Table 7), which indicates the better tendency of these compound to acquire electron density relative to amoxicillin [46].

**Table 7:** The CDFT of the best performing phenolics at the actv and allo site of PBP2x using DFT calculated by B3LYP/6-31G + (dp)

| CDFT descriptors (eV) | Amoxicillin | Silicristin | ECMG |
| --- | --- | --- | --- |
| E_LUMO | -1.0116 | -1.8273 | -1.4335 |
| E_HOMO | -6.3350 | -5.8784 | -5.9723 |
| Energy gap ($\Delta E$) | 5.3234 | 4.0510 | 4.5388 |
| Ionization energy ($I$) | 6.3350 | 5.8784 | 5.9723 |
| Electron affinity ($A$) | 1.0115 | 1.8273 | 1.4335 |
| Hardness ($\eta$) | 2.6617 | 2.0255 | 2.2694 |
| Softness ($S$) | 0.3756 | 0.4937 | 0.4406 |
| Electronegativity ($\chi$) | 3.6733 | 3.8528 | 3.7029 |
| Chemical potential ($\mu$) | -3.6733 | -3.8529 | -3.7029 |
| Electrophilicity index | 2.5346 | 3.6644 | 3.0210 |

## 3.0 Conclusion

The findings from this study revealed the diverse capability of phenolics in the binding of the actv and allo sites of PBP2x, with ECMG, silicristin, and lysidicichin having advantages in the binding of the PBP2x actv site. On the other hand, ECBE, GCG and silicristin had better affinity at the allo site. The higher affinity of silicristin at both the actv and allo sites of PBP2x showed that the compound had the best broad-spectrum effects and thus had an advantage over other top hit phenolics and amoxicillin as inhibitors of PBP2x. However, relative to other top hit phenolics, silicristin elicited higher thermodynamic entropy following binding at the actv site. This discovery might point to the necessity for further structural modifications to the molecule for stability, which could greatly boost its affinity for PBP2x. Allosteric modulation of Ser337, the critical amino acid at the actv site of PBP2x, was noted as ECBE, GCG, and silicristin with higher negative ΔGbind at the allo site instigated more fluctuation of Ser337 than amoxicillin, lysidicichin, and ECMG with moderately high affinity. This observation could imply that higher affinity of the top-hits at the allo site of PBP2x causes less obstruction of the actv site due to the lesser involvement of Ser337 in intramolecular binding of adjacent residues. Comparably, binding of the top hit phenolics and amoxicillin at the actv site of PBP2x causes less fluctuation of Ser337, suggesting that the residue could be involved in inter- and intra-molecular interaction. These findings hint to the potential advantages of top-hit phenolics, particularly silicristin, in the direct and synergistic treatment of *S. pneumoniae* infections with conventional beta-lactam medicines. More importantly, the top-hits phenolics had drug-like pharmacokinetics and toxicity profile that potentiate them as probable drug candidates. However, further *in vitro* and *in vivo* validation of the activities of the top-hits phenolics is highly suggested, and work is ongoing in this regard.

## 4.0 Materials and Methods

### 4 1. PBP2x acquisition, optimization, and identification of actv and allo sites

The experimental crystal structures of PBP2x from *S. pneumoniae* (1QMF for the actv site evaluation and 5OIZ for the allo site) were obtained from the protein data bank (PDB) (https://www.rcsb.org) [10, 12]. The crystal structures were optimized by eliminating water molecules and nonstandard amino acids using the UCSF Chimera v1.15 software tool [47]. Afterword, the x-y-z coordinates and amino acid residues at the actv {center [x (116.22), y (60.78), z (71.98], dimension [x (22.75), y (22.06), z (24.82)]} and allo site {center [x (99.33), y (67.22), z (52.41], dimension [x (21.94), y (27.41), z (20.75)]} of the protein were defined as previously reported using Discovery Studio version 21.1.0 [48] and thereafter validated from literature [10, 12].

### 4.2 Screening of phenolics using structure-based pharmacophore

Using the structure-based pharmacophore strategy specific for a ligand and a receptor [49], more than 10,000 phenolics currently known were screened against PBP2x. Briefly, a generalized pharmacophore for phenolics (Table S1) was generated from PharmaGist (https://bioinfo3d.cs.tau.ac.il/PharmaGist/php.php) using 32 phenolics. Afterwards, the generated pharmacophore was used as a query search on the ZINCPharmer database (http://zincpharmer.csb.pitt.edu) using optimized PBP2x as a receptor to aid the search. A total of 1555 phenolics (Figure S1) were obtained from the search, and the presence of the 32 phenolics used for the pharmacophore building in the total phenolics obtained served as a validation step for the analysis (Table S1).

### 4.3 Ligand acquisition, preparation, and molecular docking at the actv and allo sites of PBP2x

The library of phenolics built from the ZINCPharmer database was filtered using molecular docking techniques. Four conventional beta-lactams cefotaxime, (amoxicillin, doripenem, and aztreonam,) were used as standards for the screening. Prior to molecular docking, the Open Babel tool present on Python Prescription (PyRx) v.0.9.5 [50] was employed for the optimization of the ligands (phenolics and standards) via the addition of Gasteiger charges [34]. Subsequently, the molecular docking of the ligands was done at the actv site of PBP2x using the AutoDock Vina tool plug-in on PyRx v.0.9.5. Docking at the actv site was guaranteed via the selection of the critical residues at the actv site of PBP2x, with the grid box coordinates of the selection matching the established x-y-z coordinates (Section 2.1). After sorting the top 20 phenolics using the docking score of the reference beta-lactam antibiotics as a benchmark, the similarity of the compounds, pharmacokinetic assessment, and synthetic feasibility evaluation were used to rank the top 5 phenolics. After the top-hit phenolics acquisition from PubChem (https://pubchem.ncbi.nlm.nih.gov/) and preparation via charge addition using the UCSF Chimera v. 1.15 software program [47] the top-hit phenolics were then docked at the allo site of PBP2x using the AutoDock tool present on PyRx. The docking at the allo site was ensured by selecting the residues at the site with the coordinates of the grid box corresponding with the established x-y-z coordinates for allo site in section 4.1. For each of the top five phenolics, the docking conformation with the highest negative docking score at both the actv and allo sites of PBP2x were saved in PDB format for further molecular dynamic (MD) simulation.

To validate the docking conformation, the superimposition techniques was employed [51] where appropriate root mean square deviation (RMSD) of a docked phenolics from its reference point (points of native inhibitors) in PBP2x was calculated [51]. A relative smaller RMSD (usually < 2 Å) between the docked ligand and the native inhibitor in the response complex means comparable binding conformation [52]. The superimposition demonstrated that the top-hit phenolics, amoxicillin, and the docked native inhibitors of 1QMF (actv) and 5OIZ (allosteric) had relative binding orientation at both sites of the protein with RMSD values of 1.5 Å (actv site) and 1.4 Å (allo site). Furthermore, all the ligands interacted with the key amino acid residue in the actv (Ser337) (Figure 12a) [10] and allo (Arg426) (Figure 12b) [12] sites of PBP2x. These findings supported the validity of the docking scores obtained in the study.

**Figure 12:**
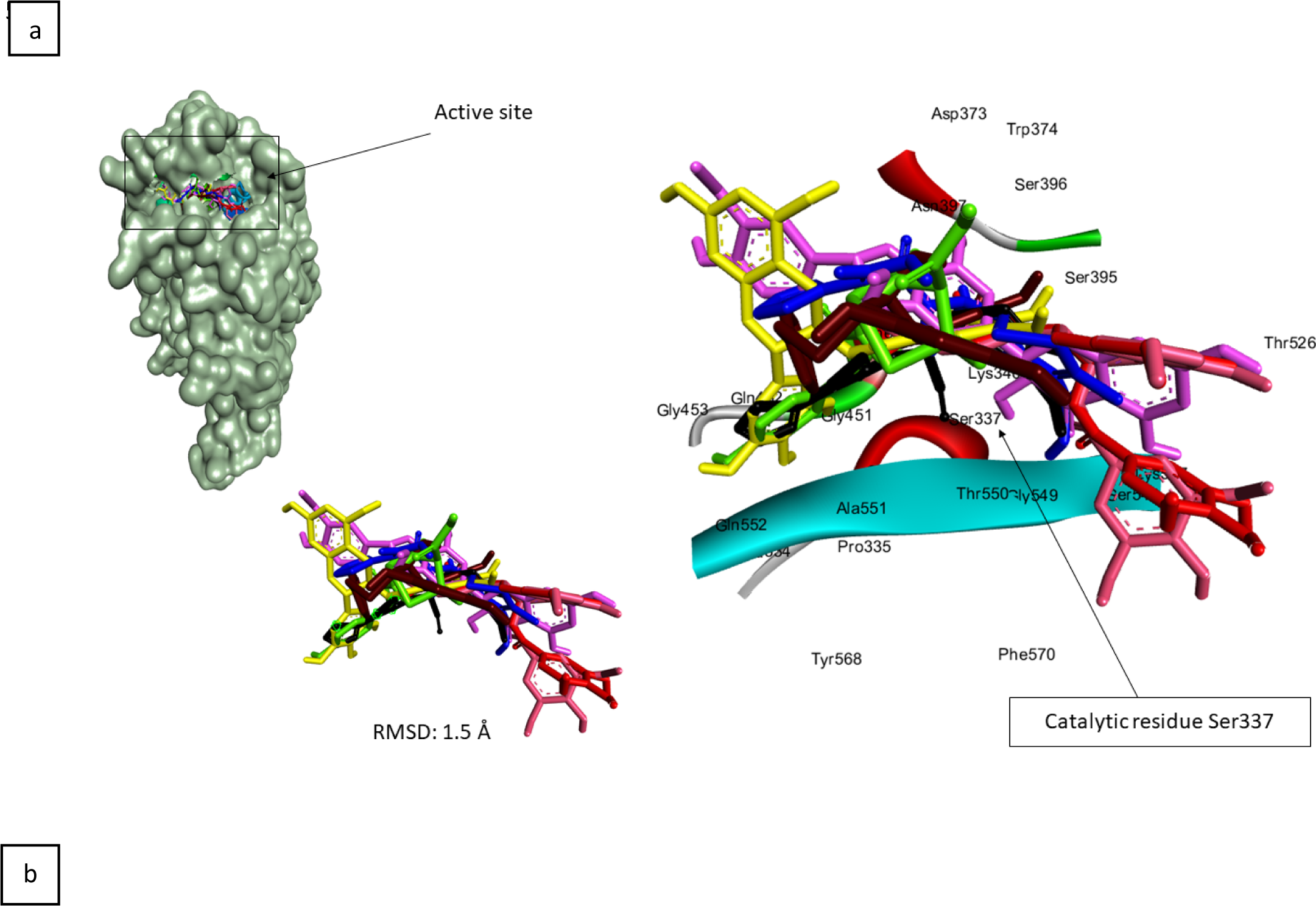
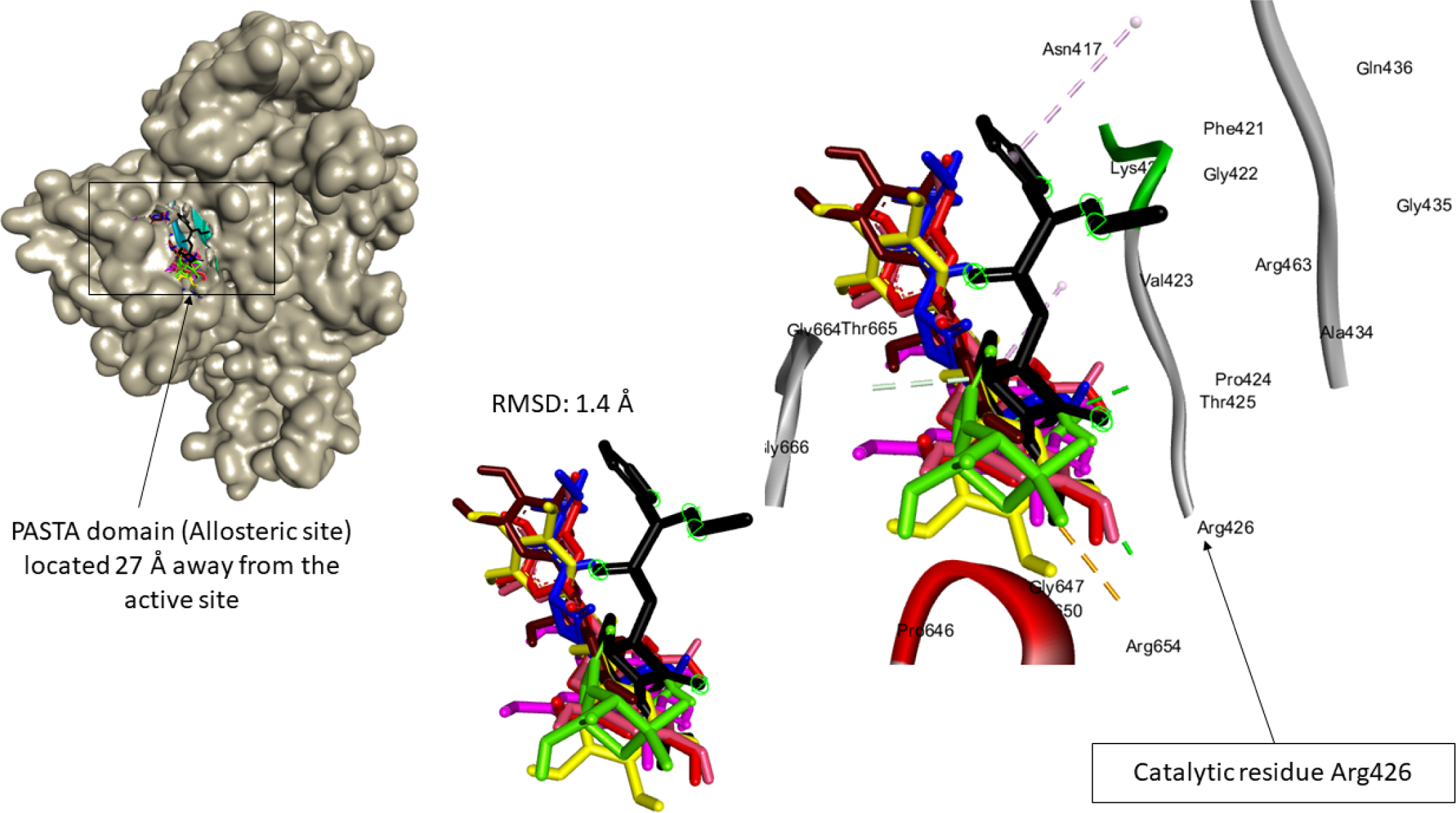
Docking protocol validation via superimposition of the docked top-hit phenolics with the experimental native inhibitor at the actv (a) and allosteric (b) sites of 1QMF and 5OIZ respectively. The docked experimental inhibitor (blue), reference standard (green) and top-hit phenolics [silicristin (magneta), ECBE (cyan), lysidicichin (red), ECMG (orange) and GCG (yellow)] all had an RMSD of 1.5 Å and 1.4 Å from the experimental native inhibitor at the actv and allo site of 1QMF and 5OIZ, respectively, suggesting partial binding orientation of the docked compounds and the experimental native inhibitors.

### 4.4. Pharmacokinetic and physicochemical prediction of phenolics

Using the simplified molecular input line entry system (SMILES)of each structure, the pharmacokinetics, physicochemical, and drug-like properties of the top twenty phenolics were predicted using the SwissADME web tools (http://swissadme.ch/index.php) and afterwards validated using the Molinspiration toolkit (https://www.molinspiration.com/cgi-bin/properties). On the other hand, the Protox II webserver (https://tox-new.charite.de/protox_II/) was employed in the prediction of the toxicity properties of the compounds. Coupled with the docking scores, the cheminformatics information obtained from these analyses were used in ranking the top hit phenolics.

### 4.5. Chemical fingerprinting of phenolics

To understand how each of the top twenty compounds is chemically similar to each other, molecular fingerprinting was carried out using Galaxy Europe (https://usegalaxy.eu./#) [53]. The information obtained from this study enables the selection of the top hit phenolics that are more diverse from each other [53]. In a nutshell, Galaxy Europe’s “molecule to fingerprint” tool was used to convert the compounds’ smile format into “Open Babel FP2 fingerprints.” The “Open Babel FP2 fingerprints” were then clustered using the fingerprinting algorithms “Taylor-Butina” and “NxN clustering,” with thresholds of 0.8 and 0.0, respectively.

### 4.6. Molecular dynamic (MD) simulations and post simulation analysis of phenolics

The AMBER 18 package of the CHPC (Centre for High Performance Computing) was used to perform a 120 ns MD simulation of each of the top hit phenolics, amoxicillin and apo-PBP2x, utilizing the AMBER force field’s FF18SB variant to describe the operating systems [54]. Using the General Amber Force Field (GAFF) and Restrained Electrostatic Potential (RESP) approaches, the ANTECHAMBER was used to create the atomic partial charges of the ligands. The protonation states were assigned using the AMBER LEaP module, which by default helps investigate and assign the correct protonation states for each amino acid residue using Na^+^, hydrogen atoms, and Cl^−^ counter ions. The numbering of residues from 1-650 was done and dipping of the systems in a manner that allows each of the atoms to be within 8 Å of any box edge was done in an orthorhombic box of TIP3P water molecules. For each of the top hit phenolics, amoxicillin and apo-PBP2x simulation, an initial equilibrium configuration (2000 steps) was achieved by a means of 500 kcal/mol restriction potential. Afterward, two 990 steps by means of steepest descent technique and conjugate degrees were carried out respectively, followed by a 990 steps unlimited minimization using the conjugate gradient approach [54]. For 50 ps, heating simulation was done gradually from 0 K to 300 K and ensuring that a constant number of atoms and volume was maintained. The potential harmonic constraint of the solutes in the systems is 10 kcal/mol, and the collision frequency is 1.0 ps. Following that, each system was equilibrated for approximately 500 ps while the operating temperature remained constant at 300 K. Restriction of hydrogen bonds using the SHAKE method was done for each simulation and each system had a randomized seeding and a 2 fs step size (corresponding to the isobaric−isothermal ensemble (NPT)). Also, a specific temperature (300 K) and constant pressure (1 bar) were maintained during each of the simulations. A Langevin thermostat was used to keep the collision frequency at 1.0 ps and the pressure-coupling constant at 2 ps [54]. Following that, the findings of the 120 ns MD simulation were analyzed and categorized as post-dynamic data. The post simulation analysis was carried out by gathering the systems’ coordinates and trajectories using the PTRAJ module. The study of RMSD, root means square fluctuation (RMSF), radius of gyration (ROG), and solvent accessible surface area (SASA) was carried out using the CPPTRAJ module, and their plots were made using Origin v 6.0. Similarly, the binding free energy (ΔGbind) was determined using an average of 120,000 snapshots collected from a 120 ns MD simulation trajectory using the equations below, utilizing the MMGBSA technique [26]

ΔGbind = G_complex_ – (G_receptor_ + G_ligand)_ ....... 1

ΔGbind = −TS + (G_sol_ + E_gas_) ......................... 2

E_ele +_ E_int_ + E_vdw_ = E_gas_ .................................. 3

- (G_GB_ - G_SA_) = G_sol_ ....................................... 4

γSASA = G_SA_ ................................................. 5

Where Egas = the gas-phase energy, E_int_ = the internal energy, E_ele_ = coulomb energy E_vdw_ = the van der Waals energy, Gsol = solvation-free energy from polar state, G_GB_ = solvation−free energy from polar state non−polar states, S = total entropy and T= temperature.

The interaction plots of the top hit phenolics and amoxicillin against PBP2x at both actv and allo sites of the protein were visualized and analyzed with Liplot v2.2.8 (Wallace *et al*., 1995).

### 4.7 Quantum chemical calculations

The purpose of this investigation was to predict the molecular characteristics of the best phenolics at the actv and allo sites of PBP2x. The most generally used DFT/B3LYP/631G/+ (d, p) basis set [43, 44] was utilized. The Centre for High Performance Computing’s Gaussian 16 program package was employed for the analysis and the results were evaluated with GaussView 6 software V. 6.0.16. The conceptual density functional theory (CDFT) descriptors such as softness, hardness, electrophilicity index, energy gap, ionization energy, and chemical potential were subsequently estimated from the energies of frontier lowest unoccupied molecular orbital (LUMO) and highest occupied molecular orbital (HOMO) consideration the Parr and Pearson interpretation [45, 55] of DFT and Koopmans theorem [46] on the correlation of ionization potential (I) and electron affinities (E) with HOMO and LUMO energy. The following equations were used for the calculations:

[E_LUMO − E_HOMO] (eV) = Energy gap (*ΔE*) ................. 1

[*I* = − E_HOMO] (eV) = Ionization energy (*I*) ...................... 2

[*A =* − E_LUMO) = Electron affinity (*A*) ............................. 3

[*η =* (*I* − *A*)/2] (eV) = Hardness (*η*) ...................................... 4

[*S =* 1/*η*] (eV) = Softness (*S*) .................................................. 5

[*χ* = (*I* + *A*)/2] (eV) = Electronegativity (*χ*) ............................ 6

[*μ =* − *χ =* − (*I* + *A*)/2] (eV) = Chemical potential (*μ*) ............ 7

[*ω = μ*^2^/2 *η*] (eV) = Electrophilicity index (*ω*) ....................... 8

## Funding

The authors specially acknowledge the financial assistance of the Directorate of Research and Postgraduate Support, Durban University of Technology (DUT), and the South African Medical Research Council (SAMRC) under a Self-Initiated Research Grant as well as the National Research Foundation (NRF) Research Development Grant for Rated Researchers (Grant number 120433) and the Competitive Programme for Rated Researchers Support (SRUG2204193723) to S. Sabiu. The views and opinions expressed are those of the authors and do not necessarily represent the official views of the funders.

## Acknowledgments

The assistance of the National Research Foundation (NRF− TWAS Doctoral Scholarship, grant number 129950), South Africa, to Mr. J. O. Aribisala is duly and thankfully acknowledged. The Centre for High−Performance Computing (CHPC), South Africa is equally acknowledged for granting access to the computing systems used in this study.

## Conflict of Interest

None to declare.

## Author’s contribution

SS conceptualized, mobilized funds, and managed the study. SS supervised the study. JOA generated and analyzed the data. JOA and NWS wrote the manuscript. All authors read and contributed to the critical review of the manuscript for intellectual content and approved the submission for publication.

## Supplementary description

Additional data on the pharmacophore classes of the phenolics, thermodynamic data at the allo site and the 2D interactions of the top-hit phenolics against the actv and allo sites are presented in the supplementary file (Supplementary Tables S1-S3, Figure S1-S4).

## Notes

### Competing Interest Statement

The authors have declared no competing interest.

